# Optimized expansion microscopy reveals species-specific spindle microtubule organization in *Xenopus* egg extracts

**DOI:** 10.1101/2024.09.11.612005

**Authors:** Gabriel Guilloux, Maiko Kitaoka, Karel Mocaer, Claire Heichette, Laurence Duchesne, Rebecca Heald, Thierry Pecot, Romain Gibeaux

## Abstract

The spindle is a key structure in cell division as it orchestrates the accurate segregation of genetic material. While its assembly and function are well-studied, the mechanisms regulating spindle architecture remain elusive. In this study, we investigate the differences in spindle organization between *Xenopus laevis* and *Xenopus tropicalis*, leveraging expansion microscopy (ExM) to overcome the limitations of conventional imaging techniques. We optimized an ExM protocol tailored for *Xenopus* egg extract spindles, improving upon fixation, denaturation and gelation methods to achieve higher resolution imaging of spindles. Our protocol preserves spindle integrity and allows effective pre-expansion immunofluorescence. This method enabled detailed analysis of the differences in microtubule organization between the two species. *X. laevis* spindles overall exhibited a broader range of bundle sizes, while *X. tropicalis* spindles contained mostly smaller bundles. Moreover, while both species exhibited larger bundle sizes near and at the spindle center, *X. tropicalis* spindles otherwise consisted of very small bundles, and *X. laevis* spindles medium-sized bundles. By enhancing resolution and minimizing distortions and fixation artifacts, our optimized ExM approach offers new insights into spindle morphology and provides a robust tool for studying the structural intricacies of these large cellular assemblies. This work advances our understanding of spindle architecture and opens up new avenues for exploring underlying mechanisms.

**SIGNIFICANCE STATEMENT:**

- Correct spindle morphology is key to its function; however, traditional microscopy methods limit our view of spindle architecture. This study addresses the gap in resolving detailed spindle microtubule organization by using advanced imaging.
- The research utilizes Expansion Microscopy (ExM) to reveal previously unobservable details of spindle morphology in egg extracts of two *Xenopus* species (*X. laevis* and *X. tropicalis*). This approach provides unprecedented clarity on microtubule arrangement and variations in spindle architecture.
- This work establishes a new protocol for high-resolution imaging of spindle structures, offering insights into how spindle architecture is adapted in differently-sized spindles to ensure proper function.

## INTRODUCTION

During cell division, the spindle plays a key role in ensuring the faithful segregation of the genetic material from the mother to daughter cells. This highly dynamic and self-organized structure is composed of thousands of microtubules and hundreds of other different proteins (Sauer *et al*., 2005), which together form a bipolar assembly that provides the necessary forces to align chromosomes and pull them towards opposite spindle poles. Therefore, any defects in this process can lead to aneuploidy, which is associated with numerous pathologies in humans (Chunduri and Storchová, 2019). Consequently, the spindle has been extensively studied across various models, significantly advancing our understanding of spindle assembly (Helmke *et al*., 2013; Guilloux and Gibeaux, 2020). Interestingly, although its components and function are well conserved across species, spindle size and morphology vary dramatically (Crowder *et al*., 2015). The precise mechanisms by which the spindle regulates its architecture and morphometrics remain incompletely understood.

The *Xenopus* egg extract is a well-established model for studying spindle assembly. This cell-free system was initially developed to investigate nuclear envelope formation and the initiation of DNA synthesis (Lohka and Masui, 1983). However, since then, it has proven to be highly versatile, allowing the assembly of cytoskeletal structures (Geisterfer *et al*., 2021). Although extracts were originally prepared from eggs of the African clawed frog, *Xenopus laevis*, the system has been adapted to various frog species, including *X. tropicalis*, *X. borealis* and *Hymenochirus boettgeri* (Brown *et al*., 2007; Kitaoka *et al*., 2018; Miller *et al*., 2019). Notably, compared to *X. laevis*, these species show different spindle sizes and architectures. In this study, we focus on the differences between the *X. laevis* and *X. tropicalis* spindles. Indeed, compared to *X. laevis, X. tropicalis* spindles are shorter (Brown *et al*., 2007), display increased microtubule density at their poles (Helmke and Heald, 2014), and have more robust kinetochore fibers (Loughlin *et al*., 2011), making their comparison a powerful tool for elucidating the principles of spindle architecture regulation. Some species-specific differences, such as the severing activity of the katanin enzyme (Loughlin *et al*., 2011), dynamic properties of tubulins (Hirst *et al*., 2020), and the TPX2/Eg5 interaction (Helmke and Heald, 2014), have been shown to contribute to global spindle size scaling between these two *Xenopus* species. However, a detailed comparison of spindle microtubule organization at much smaller scale has yet to be conducted.

While individual microtubules have a diameter of about 25 nm and can reach several micrometers in length, the spindle they organize and densely occupy ranges from about 25 µm in length for *X. tropicalis* to about 40 µm for *X. laevis*. In this context, conventional light microscopy lacks the resolution to detail the organization of single microtubules within the spindle. Super-resolution techniques have continuously pushed the diffraction limit to significantly improve resolution (Sahl *et al*., 2017), even reaching the nanometer scale (Balzarotti *et al*., 2017). However, it requires specific equipment, fluorophores and imaging conditions. On the other hand, electron microscopy can distinguish individual microtubules even when bundled at the ultrastructural scale, and has been used to image and reconstruct spindles from various species, including the yeasts *Schizosaccharomyces pombe*, *Saccharomyces cerevisiae* and *Ashbya gossypii* (Höög *et al*., 2007; Gibeaux *et al*., 2012), the worm *Caenorhabditis elegans* (Redemann *et al*., 2017), and human HeLa cells (Kiewisz *et al*., 2022). Unfortunately, this technique is laborious and time-consuming, and sample size is greatly limiting, such that the largest reconstructed spindles mentioned above are not even half the length of the *X. laevis* spindle. Previous attempts to image the whole *X. laevis* spindle faced significant challenges, such as the segmentation and tracking of thousands of microtubules within the spindle across dozens of sections and the extremely large datasets generated (Heiligenstein, 2011). Even if electron microscopy could be used to image one *X. laevis* spindle, it would be a much greater challenge to image several across various perturbed conditions to allow comparison.

For these reasons, we turned towards expansion microscopy (ExM), which enhances spatial resolution by physically enlarging the sample, allowing it to be imaged using conventional microscopes. The original ExM protocol used a trifunctional label that included a methacryloyl group for free radical polymerization, a fluorophore for visualization, and an oligonucleotide that can hybridize to a complementary sequence attached to an affinity tag for recognizing biomolecules of interest. A swellable gel is then synthesized on the sample, incorporating the labels, followed by proteinase treatment to homogenize mechanical properties, and subsequent dialysis in water to expand the gel fourfold (Chen *et al*., 2015). Since its development, various alternative protocols have been proposed to adapt ExM depending on the sample characteristics, introducing new features that have been widely adopted in later studies. For example, proExM introduced succinimidyl ester of 6-((acryloyl) amino) hexanoic acid (AcX) to anchor biomolecules directly into the polymeric gel by specifically adding acrylamides on primary amines (Tillberg *et al*., 2016). Due to significant protein loss caused by proteolytic digestion, the MAP protocol replaced it with thermal denaturation at 95°C in the presence of sodium dodecyl sulfate (SDS) (Ku *et al*., 2016). Finally, another protocol, largely used nowadays, U-ExM, was developed to expand cellular ultrastructures using a mixture of formaldehyde and acrylamide to, like AcX, link acrylamides on primary amines to anchor biomolecules in the gel, an optimized acrylamide and sodium acrylate gel composition, and thermal denaturation (Gambarotto *et al*., 2018).

Over the years, ExM has been adapted for imaging proteins, nucleic acids (Chen *et al*., 2016), lipids (Wen *et al*., 2020), and all biomolecules (Sun *et al*., 2020). Moreover, ExM has also been successfully applied to various models and tissues, such as human kidney (Zhao *et al*., 2017), *Drosophila melanogaster* (Jiang *et al*., 2018), *Caenorhabditis elegans* (Park *et al*., 2019), *Arabidopsis thaliana* (Kao and Nodine, 2019) and fungi (Götz *et al*., 2020). Nevertheless, ExM adaptations were often necessary to account for the mechanical properties of the sample and this complexity increases when mechanical characteristics vary within the sample itself. Indeed, different cellular organelles can exhibit varying expansion factors within the same cell (Büttner *et al*., 2021). As the *Xenopus* egg extract spindles are a unique subcellular structure because of their large size and high density of cross-linked microtubules, we reevaluated ExM parameters, inspired by these different protocols. Here, we successfully adapted ExM to faithfully maintain *Xenopus* egg extract structures, and ultimately analyzed the morphometric differences between *X. laevis* and *X. tropicalis* spindles with unprecedented detail. We showed that, overall, *X. laevis* spindles exhibit a broader range of microtubule bundle sizes, while *X. tropicalis* spindles are more limited to smaller bundles. We moreover revealed that, while both species favor similar larger bundle sizes near and at the spindle center, *X. tropicalis* spindles are typically built from very small bundles, and *X. laevis* spindles from medium-sized bundles in spindle regions between the poles and the midzone.

## RESULTS

### Validation of the ExM protocol on *Xenopus* egg extract spindles

We assembled spindles in *Xenopus* egg extracts supplemented with rhodamine-labeled tubulin (Figure 1A), fixed and spun them down onto coverslips (Figure 1B) as previously described (Hannak and Heald, 2006). To compare the same spindles before and after expansion and validate the tested conditions, we developed a homemade holed metal slide. This slide allowed us to image the coverslip and then recover it for ExM (Figure 1C). It was also essential to calculate the expansion factor for each spindle individually, the so-called local expansion factor, rather than assessing a global expansion factor by measuring the size of the gel before and after expansion, what was previously determined as the best practice (Truckenbrodt *et al*., 2019). Importantly, we optimized our protocol to preserve the rhodamine signal of tubulins to directly and faithfully image microtubules without the need for immunofluorescence. In addition, we aimed to retain the post-centrifugation fixation which enables pre-gelation immunofluorescence of other proteins and thus increases the options for the immunodetection of proteins of interest. To incorporate biomolecules of interest into the gel, AcX was chosen for its specificity, efficiency, and simplicity as well as to limit the use of aldehydes and prevent potential unwanted side reactions. We then began our optimization by comparing the gelation solutions from the original ExM protocol and the one from the U-ExM that uses the same components but in different proportions (Figure 1C). After polymerization, two homogenization methods were also tested: proteinase K digestion as in the ExM protocol or 95°C denaturation in the presence of SDS as in the MAP (Ku *et al*., 2016) and U-ExM protocols (Figure 1C).

**FIGURE 1:**
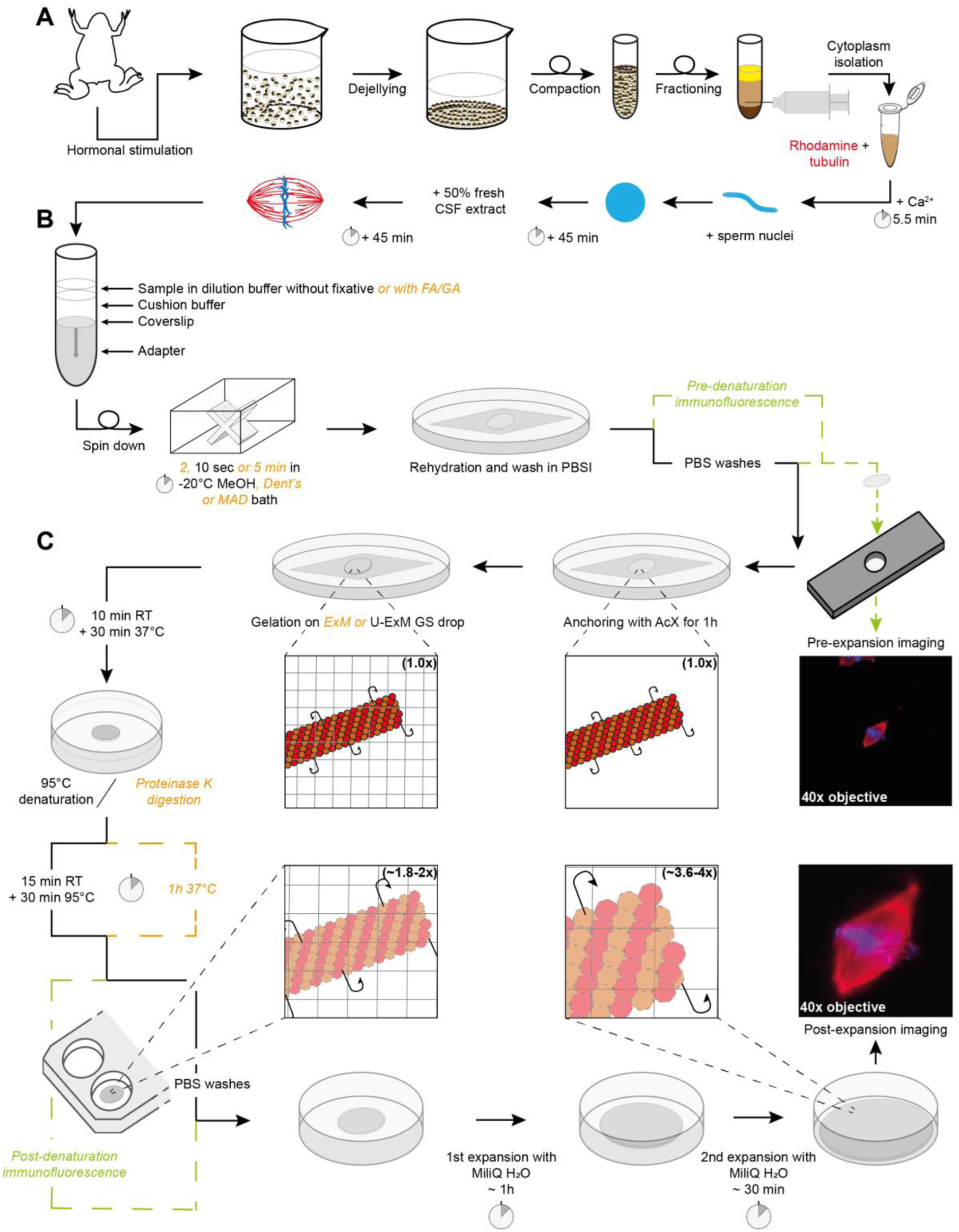
Sample processing workflow for Expansion Microscopy of *Xenopus* egg extract spindles. (A) Preparation of egg extract spindles for expansion microscopy, including major steps of egg extract preparation and spindle assembly. (B) Sample in dilution buffer, with or without fixatives, is deposited on cushion buffer to spin-down spindles onto a coverslip placed at the top of the adapter. Following the centrifugation, the coverslip is immersed inside a bath of methanol or other fixative mixtures from 2 s to 5 min, and finally rehydrated in PBSI. (C) Expansion microscopy-specific steps from anchoring and gelation to disruption and expansion. Right images show the same spindle before (top) and after (bottom) expansion with rhodamine-labeled tubulin in red and DNA in blue. Note that immunofluorescence for pre-expansion imaging is performed after sample fixation and prior to anchoring and that immunofluorescence for post-expansion imaging is done after disruption and prior to expansion for imaging. If these steps are not performed, they are replaced by PBS washes. (CSF: cytostatic factor-arrested; FA: Formaldehyde; MeOH: Methanol; Dent’s: mix of methanol and DMSO; MAD: mix of methanol, acetone and DMSO; PBSI: PBS 0.1% IGEPAL® CA-630; AcX: 6-((acryloyl) amino) hexanoic acid, succinimidyl ester; ExM: Expansion Microscopy; U-ExM: Ultrastructure ExM; RT: room temperature).

To assess the above-mentioned variations, we quantitatively compared four conditions: ExM gel with proteinase K digestion (Figure 2A), U-ExM gel with thermal denaturation (Figure 2B), ExM gel with thermal denaturation (Figure 2C), and U-ExM gel with proteinase K digestion (Figure 2D). Expansion factors were measured and their variability assessed by aligning the same spindles before and after expansion. It was first observed that proteolytic digestion (Figure 2A and 2D) induces slightly greater expansion but more variability compared to denaturation (Figure 2B-C). Additionally, for the same disruption procedure, U-ExM gels displayed slightly higher expansion factors compared to ExM gels.

**FIGURE 2:**
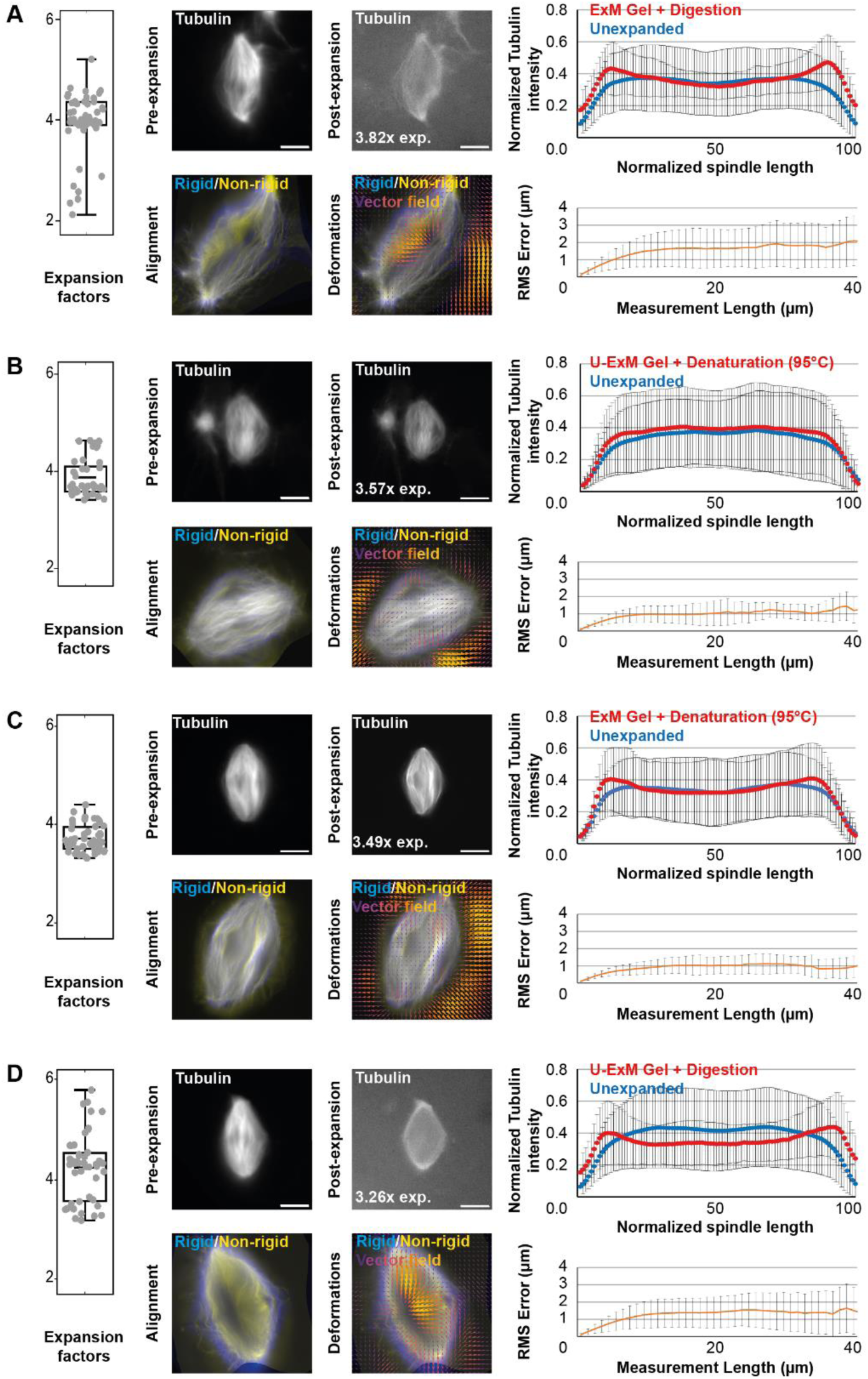
Comparison of expansion factors, rhodamine-labeled tubulin intensity distribution and RMS error between the different expansion protocols. The four tested conditions differing in the acrylamide composition of the gels and the nature of the disruption method are shown: ExM gel + proteinase K digestion (A), U-ExM gel + thermal denaturation (B), ExM gel + thermal denaturation (C), and U-ExM gel + proteinase K digestion (D). Left panels are boxplots showing individual spindle expansion factors (n = 51, n = 41, n = 47, n = 47 spindles from 4 egg extracts respectively for A, B, C, and D), the center line is the mean, the box indicates the 25%-75% quantiles and the error bars the minimum and maximum values. Average expansion factors were A: 3.94 ± 0.63, B: 3.86 ± 0.37, C: 3.71 ± 0.27, D: 4.22 ± 0.67 (average value ± standard deviation). For each, top images are representative images of unexpanded (left) and expanded (right) spindles, with rhodamine-labeled tubulin shown in gray. The expansion factor of the shown spindle is indicated. On the right of the top images are the average normalized fluorescence intensity distributions ± standard deviation for unexpanded (blue) and expanded (red) spindles. Bottom images show the alignment (left) made between rigid registration and non-rigid registration, and (right) the vector field of deformations calculated from the registrations. On the right of the bottom images, panels represent the Root-Mean-Square length measurement error as a function of measurement length for pre- vs. post-expansion images. The orange line shows the average and the error bars, the standard deviation (n = 51, n = 41, n = 47, n = 47 spindles from 4 egg extracts, respectively for A, B, C, and D). Scale bars, 10 μm for unexpanded spindles and 40 µm for expanded spindles.

Next, the preservation of rhodamine-labeled tubulin fluorescence distribution was analyzed by measuring tubulin intensity along the axis of the spindle from pole to pole in both pre- and post-expansion spindles (top images for each condition in Figure 2). It appeared evident that proteolytic digestion resulted in decreased intensity in the center of the expanded spindle (blue and red curves in Figure 2A and 2D), and to a lesser extent, in the ExM gel with thermal denaturation condition (blue and red curves in Figure 2C). In contrast, the U-ExM gel with thermal denaturation condition (blue and red curves in Figure 2B) showed a similar distribution of tubulin across both unexpanded and expanded spindles.

Finally, to assess possible distortions due to expansion, we calculated the RMS error as described in the original ExM protocol (Chen *et al*., 2015). The unexpanded spindle images were magnified fourfold and rigidly aligned by rotating and resizing them to match the expanded spindle images using the Elastix software. Non-rigid alignment with allowed local deformations was then performed to perfectly fit the two images. For each alignment, the applied transformations were registered and applied to expanded spindles to generate two new images, one having undergone rigid transformations, and the other having undergone rigid and non-rigid transformations. These two images were aligned in Mathematica (bottom left images for each condition in Figure 2) and the differences between the two images (representing the deformations due to expansion) were calculated in a vector field (bottom right images for each condition in Figure 2) and given as RMS error length measurement (orange curves in Figure 2). Doing so, we observed that proteolytic digestion resulted in higher RMS errors compared to thermal denaturation, indicating more distortions.

Based on these observations, we decided to proceed with the U-ExM gel and thermal denaturation (Figure 2B) as it provided significant expansion with minimal variations, preservation of the rhodamine-labeled tubulin signal across the spindle, and acceptable distortions, with a 1 µm measurement error on 40 µm length, representing a 2.5% distortion error, which is within most published protocols’ range of 1 to 5% (Chen *et al*., 2015; Ku *et al*., 2016; Tillberg *et al*., 2016; Gambarotto *et al*., 2018).

### Expansion successfully increases resolution

To assess the increased resolution, the same spindles were imaged before and after expansion with confocal microscopy equipped with a gain of resolution technology. These spindles were first observed in three dimensions using Napari software to visualize the overall gain of resolution. Individual microtubule bundles could easily be distinguished within spindles and the DNA was correctly expanded (Figure 3A). To demonstrate this increased resolution, we focused on spindle pole regions, where single microtubules were visible, and plotted the fluorescence distribution across microtubule bundles and observed that microtubules were more clearly resolved after expansion (Figure 3B). However, microtubules appeared buckled within the spindles. To confirm that this was not due to our expansion protocol, we compared the undulations in unexpanded and expanded spindles (Figure 3C). As microtubules were already buckled before expansion, we concluded that this was likely due to the fixation protocol. Indeed, aldehydes present in our dilution buffer are known to induce wavy microtubules (Cross and Williams Jr., 1991), and methanol baths used can cause cell shrinkage (Melan, 1999), potentially similarly shrinking spindles and thus causing microtubule buckling.

**FIGURE 3:**
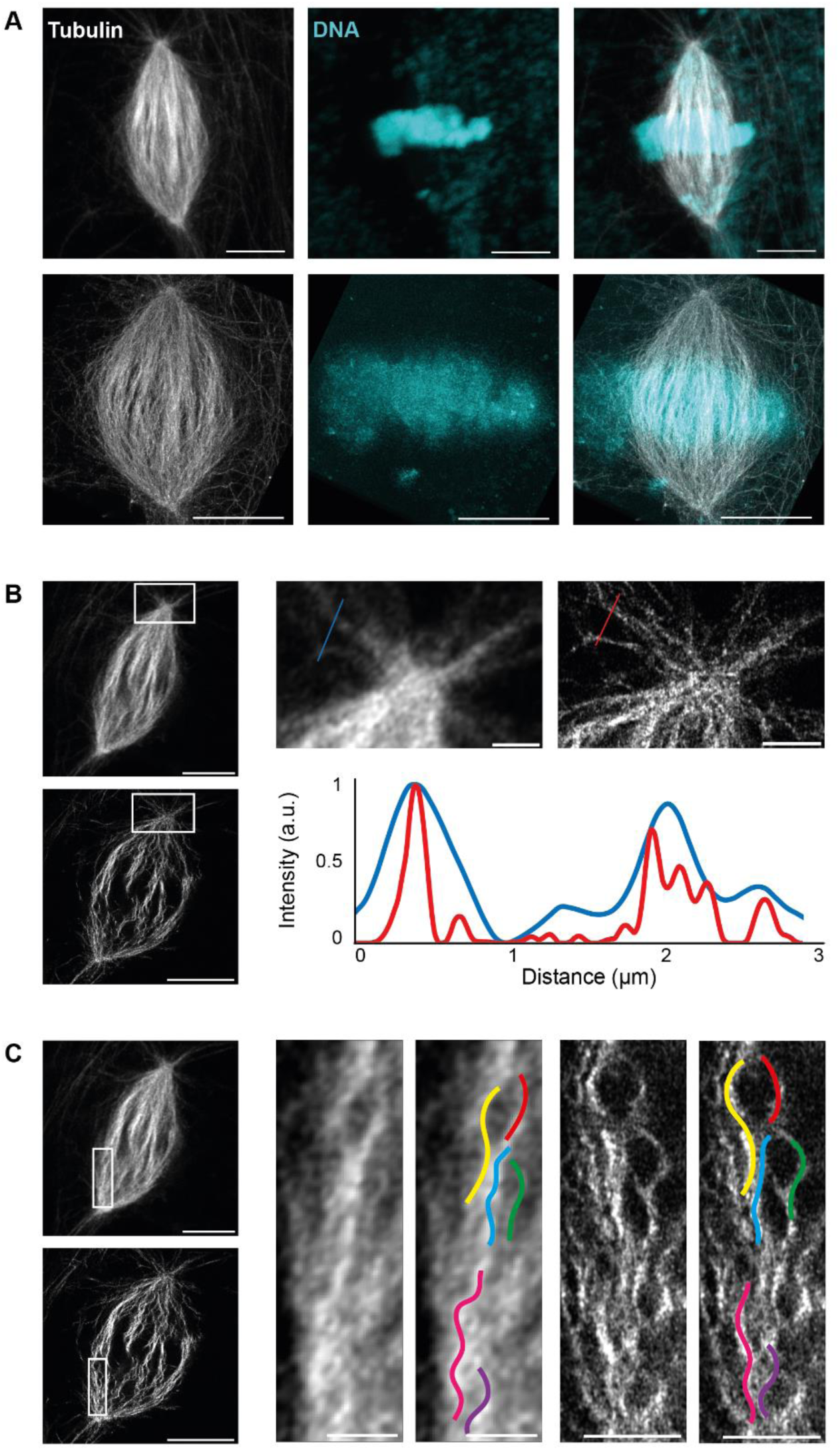
Analysis of resolution improvement and fixation artifacts. (A) Images from a 3 dimension rendering in Napari showing an unexpanded (top) and expanded (bottom) spindle with rhodamine-labeled tubulin signal (left), DNA signal (middle), and merged signals (right). (B) Single z plane of the spindle shown in (A), unexpanded (top) and expanded (bottom). Magnified views of boxed regions (top right). Profiles of tubulin intensity along the blue and red lines (bottom right). (C) Single z plane of the spindle shown in (A), unexpanded (top) and expanded (bottom). Magnified views of boxed regions with colored curves traced on unexpanded (left) and expanded (right) microtubule bundles. Scale bars, 10 µm for unexpanded spindles and 40 µm for expanded spindles in A, B and C, 2 µm for magnified views of boxed regions showing unexpanded microtubules and 8 µm for expanded microtubules in B and C.

Therefore, we attempted to improve the classical spin-down protocol by modifying various parameters. To keep post-centrifugation fixation to enable pre-expansion immunofluorescence, we began by changing the formaldehyde incubation duration, using dilution buffer with or without detergent, changing the post-methanol rehydration agent, and reducing the centrifugation speed (Supplemental Figure S1). We also varied the formaldehyde concentration, removing it, or replacing it with glutaraldehyde at different concentrations (Supplemental Figure S2A and B). While fixing spindles with glutaraldehyde seemed promising before expansion, with straighter microtubules, these were deformed and appeared crushed after expansion (Supplemental Figure S2C), displaying increased distortions with more than 10% error (Supplemental Figure S2D). Moreover, stronger fixation resulted in less successful incorporation within the gel. Altogether, these attempts led us to decide to remove any fixative from the dilution buffer in our final conditions and supported our choice of AcX for anchoring egg extract spindles.

### Minimal post-centrifugation fixation better preserves spindle integrity

In parallel to pre-centrifugation fixation optimization tests, we optimized the post-centrifugation methanol fixation. We tested alternatives, including Dent’s fixative (80% methanol, 20% DMSO; (Dent *et al*., 1989)), MAD (40% methanol, 40% acetone, 20% DMSO; (Gibeaux *et al*., 2018)), and a variant of the latter, MAD2 (60% methanol, 20% acetone, 20% DMSO), with the objective of tempering the methanol dehydration (Figure 4). Visually, the MAD and Dent’s alternatives appeared superior in terms of microtubule preservation, with more individualized bundles. Importantly, both the tubulin distribution intensity along spindles and RMS errors were similar across all conditions (Figure 4A), suggesting that the expansion was not affected by these modifications. Only the expansion factors of samples fixed with alternative baths were significantly higher compared to the control (Figure 4B).

**FIGURE 4:**
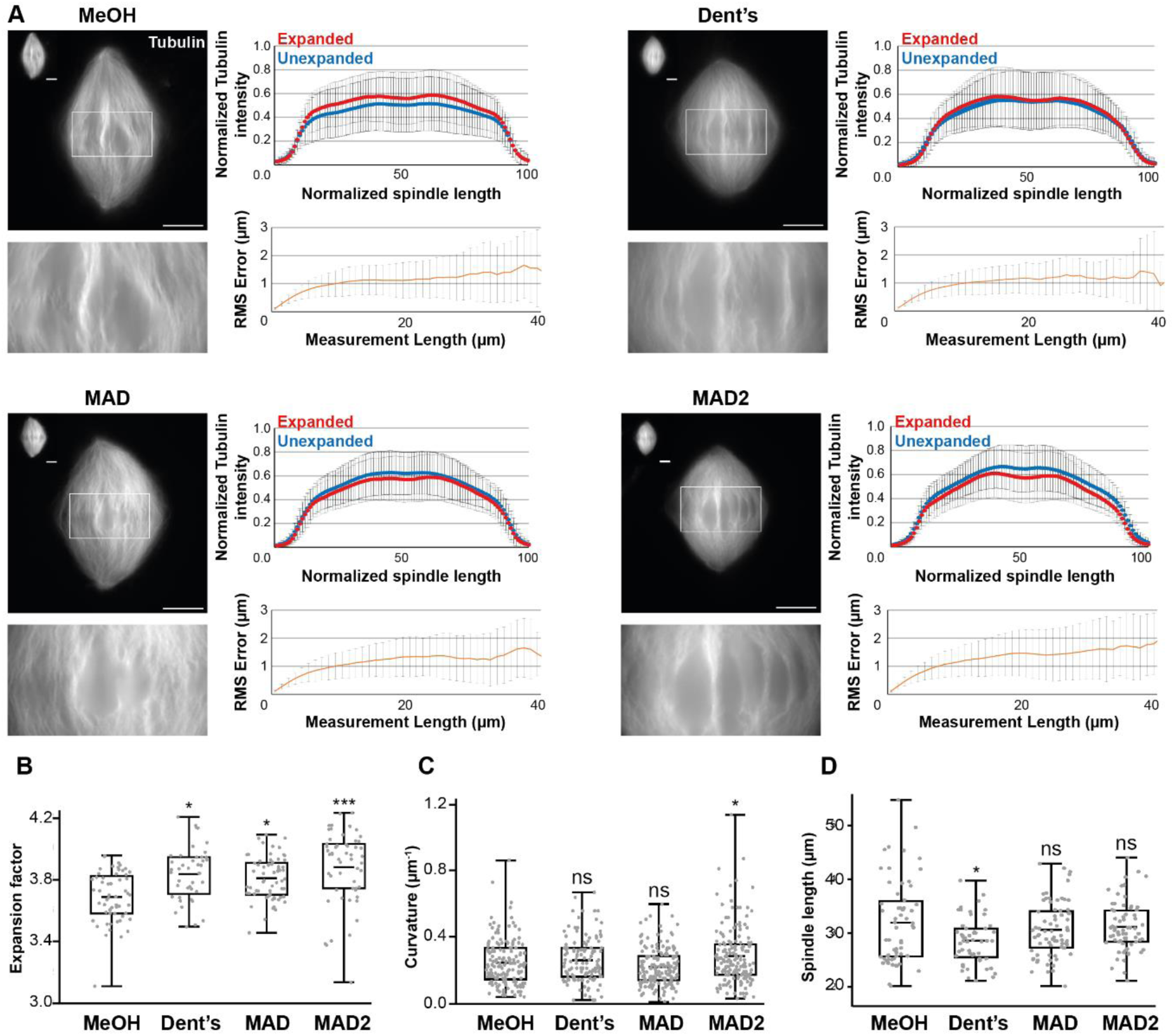
Evaluating the effects of changing the nature of the post-centrifugation fixation. (A) The four tested conditions differing by the composition of the post-centrifugation fixation: MeOH (100% methanol), Dent’s (80% methanol, 20% DMSO), MAD (40% methanol, 40% acetone, 20% DMSO), MAD2 (60% methanol, 20% acetone, 20% DMSO). For each top left images show expanded spindles (rhodamine-labeled tubulin signal) and their corresponding unexpanded spindles in the top left corner of the image. Bottom left images are magnified views of boxed regions in top images. Top right panels are the average normalized fluorescence intensity distributions ± standard deviation for unexpanded (blue) and expanded (red) (n= 57 spindles from 6 egg extracts, n = 47 spindles from 5 egg extracts, n= 64 from 5 egg extracts, n = 60 spindles from 6 egg extracts, respectively for MeOH, Dent’s, MAD and MAD2). Bottom right panels represent the Root-Mean-Square length measurement error as a function of measurement length for pre- vs. post-expansion images. The orange line shows the average and the error bars, the standard deviation (n = 56 spindles from 6 egg extracts, n = 44 spindles from 5 egg extracts, n = 64 spindles from 5 egg extracts, n = 54 spindles from 6 egg extracts, respectively for MeOH, Dent’s, MAD and MAD2). (B) Boxplots showing individual spindle expansion factors (same statistics as for Root-Mean-Square error study), the center line is the mean, the box indicates the 25%-75% quantiles and the error bars the minimum and maximum values. Average expansion factors were MeOH: 3.68 ± 0.16, Dent’s: 3.83 ± 0.17, MAD: 3.80 ± 0.14, MAD2: 3.88 ± 0.23 (average value ± standard deviation). (C) Boxplots showing average curvature of microtubules measured by tracking individual microtubule bundles on expanded spindles (n = 159 microtubules, n = 122 microtubules, n = 173 microtubules, n = 168 microtubules, respectively for MeOH, Dent’s, MAD and MAD2), the center line is the mean, the box indicates the 25%-75% quantiles and the error bars the minimum and maximum values. Average curvature were MeOH: 0.25 µm^-1^ ± 0.14, Dent’s: 0.27 µm^-1^ ± 0.13, MAD: 0.23 µm^-1^ ± 0.11, MAD: 0.29 µm^-1^ ± 0.17 (average value ± standard deviation). (D) Boxplots showing individual spindle length measured on unexpanded spindles (n = 59 spindles from 6 egg extracts, n = 52 spindles from 5 egg extracts, n = 61 spindles from 5 egg extracts, n = 61 spindles from 6 egg extracts, respectively for MeOH, Dent’s, MAD and MAD2), the center line is the mean, the box indicates the 25%-75% quantiles and the error bars the minimum and maximum values. Average spindle length were MeOH: 31.93 µm ± 7.72, Dent’s: 28.57 µm ± 4.26, MAD: 30.62 µm ± 4.99, MAD2: 31.12 µm ± 4.53 (average value ± standard deviation). Scale bars, 10 μm for unexpanded spindles and 40 µm for expanded spindles. * *p* < 0.05, ** *p* < 0.01, *** *p* < 0.001, *ns* non-significant.

Since microtubule buckling was the most evident indicator of protocol issues, microtubule average curvature was measured using the Fiji plugin Kappa. Although microtubules seemed less curved with less variability for MAD bath (0.23 µm^-1^ ± 0.11 vs. 0.25 µm-1 ± 0.14; average ± standard deviation), consistent with more individualized bundles, no improvement was statistically significant at this level (Figure 4C). The other parameter influenced by chemical fixation is shrinkage of the fixed structures. We therefore measured unexpanded spindle lengths in each condition but did not see any significant differences, except from an even greater reduction in size from Dent’s fixative condition (Figure 4D). Together, although the MAD fixation seemed to maintain better individualized microtubule bundles, these results do not indicate a clear improvement of the fixation protocol. However, as the MAD mix aimed to temper the strong dehydration and extraction effect of methanol, we wondered if we could get significantly better results by reducing the methanol fixation duration.

To test this idea, we reduced the duration of both control and MAD fixation and from 5 min to 2 s. Additionally, from these new conditions on, based on our previous results (Supplemental Figure S1 and Supplemental Figure S2), we removed the fixative from the dilution buffer and directly assessed the influence of post-centrifugation baths. Visually, microtubules appeared straighter and more individualized with reduced fixation durations. Of note, reduced durations also led to the disappearance of spindles between pre-expansion to post-expansion imaging, likely due to them washing off the coverslip during washes, which was most striking in the 2 s condition.

We measured the distribution of rhodamine-labeled tubulin intensity along the spindles. For both methanol 10 s and MAD 10 s conditions (Figure 5A), the overlap was nearly perfect. Yet, we observed an increase in distortions as fixation duration decreased, with the RMS error approaching 2 µm in the methanol 10 s condition and almost 3 µm in the MAD 10 s condition (Figure 5A), and more than 3 µm in the 2 s conditions (Figure 5A). Interestingly, the methanol 10 s condition showed an increased expansion factor while it decreased in its MAD counterpart (Figure 5B).

**FIGURE 5:**
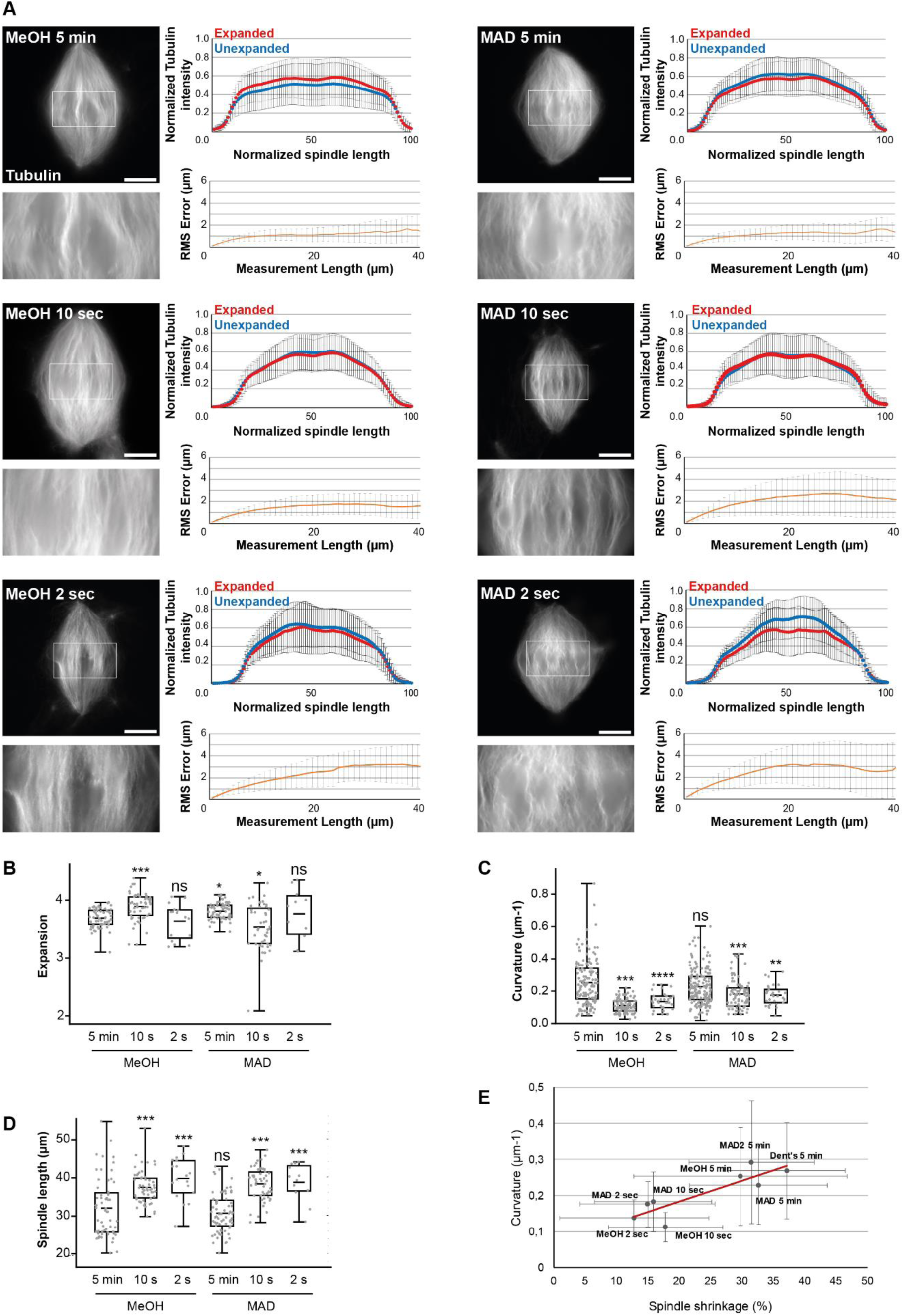
Evaluating the effects of decreasing the duration of the post-centrifugation fixation. (A) The six tested conditions differing by the composition and duration of the post-centrifugation fixation: MeOH (100% methanol, left), MAD (40% methanol, 40% acetone, 20% DMSO, right), the durations are indicated on each spindle image. Each top left images show expanded spindles (rhodamine-labeled tubulin signal). Corresponding bottom left images are magnified views of boxed regions in top images. Top right panels are the average normalized fluorescence intensity of tubulin distributions ± standard deviation for unexpanded (blue) and expanded (red) (n= 57 spindles from 6 egg extracts, n = 51 spindles from 4 egg extracts, n= 17 from two egg extracts, n = 64 spindles from 5 egg extracts, n= 45 spindles from 4 egg extracts, n = 11 spindles from two egg extracts respectively for MeOH 5 min, MeOH 10 s, MeOH 2 s, MAD 5 min, MAD 10 s and MAD 2 s). Bottom right panels represent the Root-Mean-Square length measurement error as a function of measurement length for pre- vs. post-expansion images. The orange line shows the average and the error bars, the standard deviation (n= 56 spindles from 6 egg extracts, n = 52 spindles from 4 egg extracts, n= 16 spindles from two egg extracts, n = 64 spindles from 5 egg extracts, n= 44 from 4 egg extracts, n = 11 spindles from two egg extracts respectively for MeOH 5 min, MeOH 10 s, MeOH 2 s, MAD 5 min, MAD 10 s and MAD 2 s). (B) Boxplots showing individual spindle expansion factors (same statistics as for Root-Mean-Square error study), the center line is the mean, the box indicates the 25%-75% quantiles and the error bars the minimum and maximum values. Average expansion factors were MeOH 5 min: 3.68 ± 0.16, MeOH 10 s: 3.88 ± 0.23, MeOH 2 s: 3.63 ± 0.30, MAD 5 min: 3.80 ± 0.14, MAD 10 s: 3,53 ± 0,39, MAD 2 s: 3,76 ± 0,43 (average value ± standard deviation). (C) Boxplots showing average curvature of microtubules measured by tracking individual microtubule bundles on expanded spindles (n = 159 microtubules, n = 112 microtubules, n = 34 microtubules, n = 173 microtubules, n = 90 microtubules, n = 30 respectively for MeOH 5 min, MeOH 10 s, MeOH 2 s, MAD 5 min, MAD 10 s and MAD 2 s), the center line is the mean, the box indicates the 25%-75% quantiles and the error bars the minimum and maximum values. Average curvature were MeOH 5 min: 0.25 µm^-1^ ± 0.14, MeOH 10 s: 0.11 µm^-1^ ± 0.04, MeOH 2 s: 0.14 µm^-1^ ± 0.05, MAD 5 min: 0.23 µm^-1^ ± 0.11, MAD 10 s: 0.18 µm^-1^ ± 0.08, MAD 2 s: 0.18 µm-^1^ ± 0.06 (average value ± standard deviation). (D) Boxplots showing individual spindle length measured on unexpanded spindles (n= 56 spindles from 6 egg extracts, n = 58 spindles from 4 egg extracts, n= 18 from two egg extracts, n = 72 spindles from 5 egg extracts, n= 50 spindles from 4 egg extracts, n = 12 spindles from two egg extracts respectively for MeOH 5 min, MeOH 10 s, MeOH 2 s, MAD 5 min, MAD 10 s and MAD 2 s), the center line is the mean, the box indicates the 25%-75% quantiles and the error bars the minimum and maximum values. Average spindle length were MeOH 5 min: 31.93 µm ± 7.72, MeOH 10 s: 37.35 µm ± 4.14, MeOH 2 s: 39.63 µm ± 5.39, MAD 5 min: 30.62 µm ± 5.00, MAD 10 s: 38.24 ± 4.25, MAD 2 s: 38.65 ± 4.88 (average value ± standard deviation). (E) Representation of average curvature of microtubules as a function of spindle shrinkage. Black dots, values for indicated conditions, red line, linear trend curve (R² = 0.74). Scale bars, 40 µm. * *p* < 0.05, ** *p* < 0.01, *** *p* < 0.001, *ns* non-significant.

Importantly, the shorter duration conditions resulted in less curved microtubules (Figure 5C), and longer spindle lengths (Figure 5D). We found a positive correlation between average spindle microtubule curvature and average spindle shrinkage (Figure 5E; R² = 0.74), suggesting that spindle shrinking resulting from chemical fixations led to microtubule buckling.

Taken together, post-centrifugation fixation with methanol for 10 s gave the best compromise, (i) sufficiently fixing the spindles onto the coverslip with (ii) moderate spindle shrinkage, (iii) reasonable distortion error below 5%, which remains within the range of published protocols, (iv) the straightest microtubules of all conditions and (v) a perfect maintenance of the rhodamine-labeled tubulin fluorescence distribution. We therefore selected this fixation condition as our final fixation protocol, in which chemical fixation (no aldehydes and a short dehydration step) was effective and minimally deleterious.

One goal of our optimization was to enable pre-expansion immunofluorescence. Indeed, while post-expansion immunofluorescence can be performed with our protocol as with any other ExM protocol, some antibodies are not functional after expansion, likely as a result of the denaturation step. This was the case for most antibodies routinely used in the lab for classical immunofluorescence analysis in *Xenopus* egg extract. To confirm that pre-expansion immunofluorescence was still possible after 10 s of methanol fixation, we performed immunofluorescence using an anti-dynein antibody that displays a faint signal in classical immunofluorescence analysis and does not work for post-expansion immunofluorescence. (Supplemental Figure S3A). As expected with this antibody, a faint signal on the spindle with increased signal at the spindle poles was observed. After retrieving the coverslip, we applied our expansion protocol and imaged the gel using an epifluorescence microscope (Supplemental Figure S3B). The labeling was properly preserved after expansion, together with well separated microtubule bundles. Moreover, the increased signal at the poles observed pre-expansion appeared as separated puncta and few separated dots become visible within the spindle after expansion. These findings confirm that our protocol allows pre-expansion immunofluorescence and suitable for antibodies that cannot be used for post-expansion immunofluorescence.

### Individual microtubule bundles can be segmented from expanded spindles

With our final protocol in hand, we returned to high-resolution imaging and observed that, compared to spindles in Figure 3A, the spindle volume was better preserved and the microtubules were incomparably straighter and individualized (Figure 6A & Supplemental Movie S1). Measurement of the effective resolution under this condition, as previously described (Chen *et al*., 2015) revealed expanded, clearly individualized, microtubules with a full-width at half-maximum (FWHM) of 40.8 ± 6.5 nm (mean ± standard deviation, n = 22 microtubules from three egg extracts). To estimate the effective resolution, we deconvolved our observed microtubule FWHM using the commonly accepted microtubule width of 25 nm (Weber *et al*., 1978), and obtained an effective resolution for our protocol of ∼32.3 nm. This is comparable to the theoretical maximum resolution of Airyscan (140 nm) divided by the expansion factor (3.88). Furthermore, we found that this protocol also perfectly preserved spindles assembled in *X. tropicalis* egg extracts (Figure 6B & Supplemental movie S2).

**FIGURE 6:**
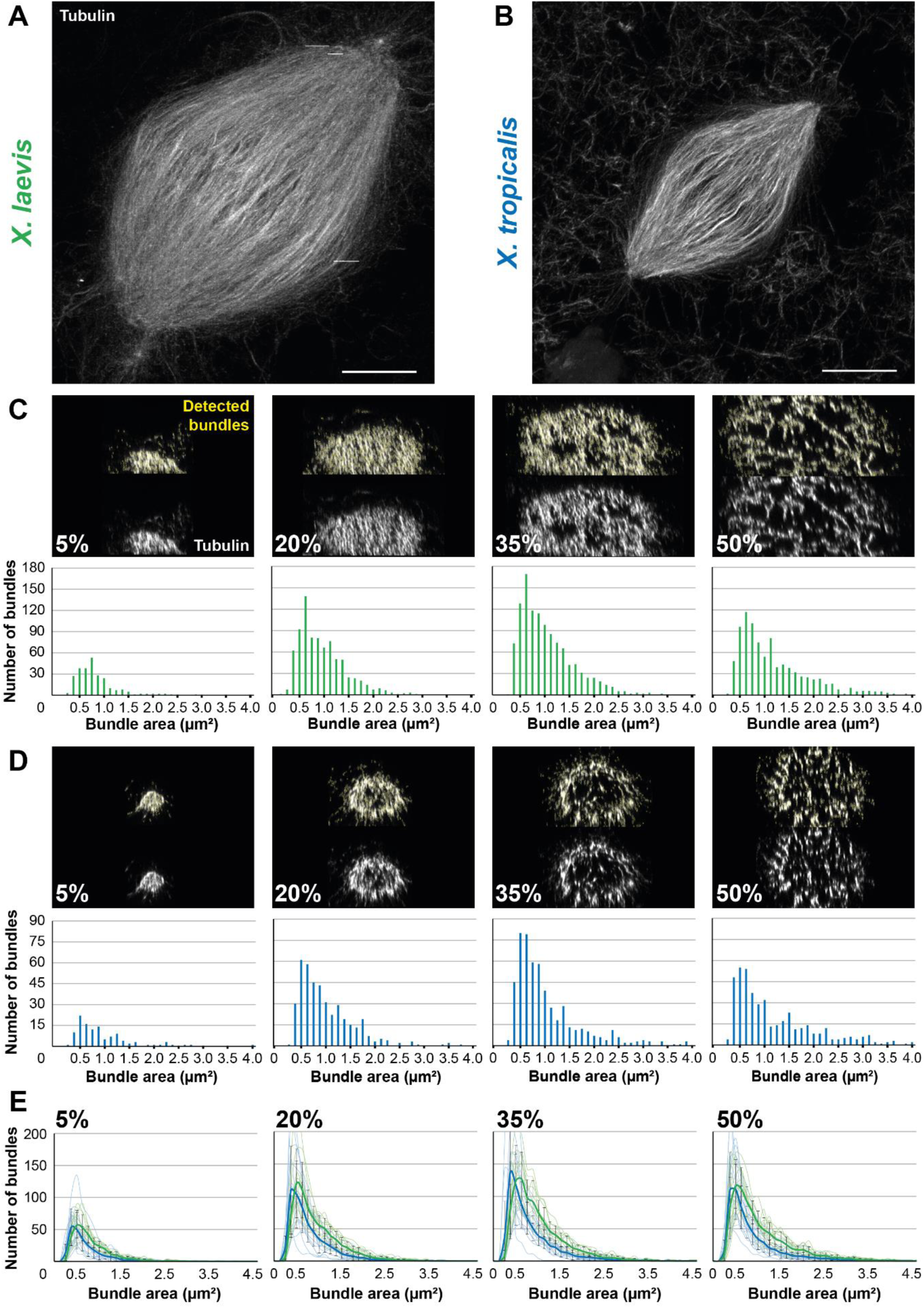
Development of an analysis workflow to analyze and compare expanded spindles from *X. laevis* and *X. tropicalis*. (A) Expanded spindle from *X. laevis* egg extract with rhodamine-labeled tubulin signal, imaged with Airyscan microscope, image taken from a 3 dimensional rendering in Napari. (B) Expanded spindle from *X. tropicalis* egg extract with rhodamine-labeled tubulin signal, imaged with Airyscan microscope, image taken from a 3 dimensions rendering in Napari. (C) Top images show cross-sectioned microtubule bundles from the spindle in (A) (respective sections are indicated in bottom left corners) before (bottom) and after (top) bundle detection (in yellow) by the ImageJ macro. Bottom bar plots show the distribution of these detected bundles, according to their area. (D) Top images show cross-sectioned microtubule bundles from the spindle in (B) (respective sections are indicated in bottom left corners) before (bottom) and after (top) bundle detection (in yellow) by the Fiji macro. Bottom bar plots show the distribution of these detected bundles according to their area. (E) Distribution of the average number of bundles according to their area ± standard deviation for spindles from *X. laevis* (green, n = 11 spindles from three egg extracts) and *X. tropicalis* (blue, n = 15 spindles from three egg extracts) in indicated cross-sections. For (C-E) bundle areas are binned into classes of 0.125 µm^2^. Each bin is named according to its upper range value. Scale bars, 20 µm.

To decipher the architectural differences between the spindles of these two species, we performed a comparative analysis at an unprecedented resolution. To do so, we developed an analysis workflow enabling us to observe the organization of microtubules from within the expanded spindles. The workflow was designed to reconstruct a spindle in three dimensions and spatially rotate it, using selected pole coordinates, to align the pole-to-pole axis with the z-axis. As a result, we were able to cross-section the spindle at various chosen positions and apply a specifically-trained Stardist model to quantify microtubule bundles in these sections, and determine their individual area and fluorescence intensity. We chose to section at the poles (5% and 95% sections), at the spindle center (50% section), and between these locations (at 20%, 35%, 65%, and 80% sections) to understand the organization of microtubules within the spindles. Since our microtubules were labeled with rhodamine-labeled tubulin, it was expected that increased microtubule number in a bundle would increase bundle size and measured fluorescence intensity. Therefore, to assess the accuracy of our bundle detection, in addition to visual inspection, we plotted the cross-section area against the intensity for each detected bundles for *X. laevis* (Supplemental Figure S4A) and *X. tropicalis* (Supplemental Figure S4B), and observed a satisfactory linear relationship between these two parameters.

We analyzed the number of microtubule bundles in each section according to their area, from spindles of *X. laevis* (Figure 6C) and *X. tropicalis* (Figure 6D). A similar distribution between the two species was observed, with a majority of bundles being 0.5-1.0 µm^2^, and a higher count for *X. laevis* as expected due to their respective spindle size. However, before further analyzing these data, we performed additional validations. We compared the distribution of fluorescence intensity, as commonly used (Kitaoka *et al*., 2018), using a 10-pixel-wide line scan on unexpanded spindles, with a size-corresponding ROI on expanded spindles between the two *Xenopus* species (Supplemental Figure S4C). We observed similar distributions between the two methods with a greater dip in intensities at the *X. tropicalis* spindle center. Conversely, we then analyzed unexpanded spindles with line scans covering the full spindle width, more consistent with our analysis workflow on expanded spindles (Supplemental figure S4D). Doing so, we observed a generally higher total intensity for *X. laevis*, which was expected given their size differences. Of note, for *X. tropicalis* spindles, which are ∼30% shorter (Brown *et al*., 2007), the maximum measured intensity in expanded spindles was approximately also 30% lower than for *X. laevis* (60 a.u. vs. 90 a.u.). Interestingly, the number of detected bundles in total spindles displayed the same global distributions between the two *Xenopus* species, with a higher number for *X. laevis* (again a difference of about 30% between the two species). The same distributions were observed for the small ROI analysis, however the counts were higher for *X. tropicalis*, suggesting that microtubules may be denser in these spindles (Supplemental figure S4C-D).

Comparing the average number of bundles according to their area between *X. laevis* (n = 11 spindles) and *X. tropicalis* (n = 15 spindles) revealed that the majority of bundles across the sections were approximately 0.5 µm^2^ in cross-section area. Considering expanded microtubules as circles of 200 nm in diameter (25 nm for the microtubule plus 25 nm for a crosslinker multiplied by a 4x expansion) to fit in circles of the bundle cross-section area, we estimated that a 0.5 µm^2^ bundle cross-sections would correspond to ∼10 microtubules, and 4 µm^2^ to ∼100 microtubules. A broader range of bundle sizes was nevertheless observed in *X. laevis* spindles, where it was more restricted to smaller sizes in *X. tropicalis* (Figure 6E). Furthermore, we observed a progressive increase in the maximum number of bundles along the spindle, from about 50 bundles in 5% section to 110 in 20%, and 130 in 35%, and decreasing to 110 in 50%. This is consistent with the shape of the spindle and the presence of chromosomes at the spindle center (Figure 6E).

Altogether, the linearity between bundle cross-sectional areas and intensities, the correspondence observed between tubulin distributions in unexpanded and expanded spindles, and the coherent distribution of bundle numbers across different spindle sections, confirmed the capacity of our analysis workflow to segment, analyze, and compare microtubule bundle organization between the two *Xenopus* species.

### Differently sized microtubule bundles are differentially localized along the spindle length between *X. laevis* and *X. tropicalis*

To analyze the distribution of microtubule bundles in further detail, we summed their areas within each area size bins (Figure 7A). First, we examined, for each bundle area bin, their summed area along the length of the spindle and compared them between the two *Xenopus* species. Interestingly, for the smaller bundles (areas of 0.25 and 0.375 µm^2^), the total areas they occupied within the entire spindle was greater for *X. tropicalis* than for *X. laevis*. Notably, their prominence was higher in the 35% and 65% sections, with a dip at the spindle center in *X. tropicalis*, while they were mostly homogeneously distributed along the spindles in *X. laevis*. The relative proportions between the two species became equal for the 0.5 µm^2^ bundles and were higher for all the larger ones in *X. laevis*. It was also observed that bundles from 0.125 to 0.375 µm² for *X. tropicalis* (exemplified with 0.25 and 0.375 µm² classes in Figure 7A) and from 0.75 to 1.375 µm² for *X. laevis* (exemplified with 1 µm² class in Figure 7A) were very strongly represented in the 35% and 65% with a dip at the metaphase plate. However, from 2.0 µm^2^ bundles, their distribution shifted to become more represented at the 50% section of both *X. laevis* and *X. tropicalis* spindles, which is even more pronounced for bigger bundles from 3 µm² (exemplified with 3 and 4 µm² classes in Figure 7A). This suggests that larger bundles, likely the more robust ones, are required at the spindle midzone to ensure spindle bipolarity.

**FIGURE 7:**
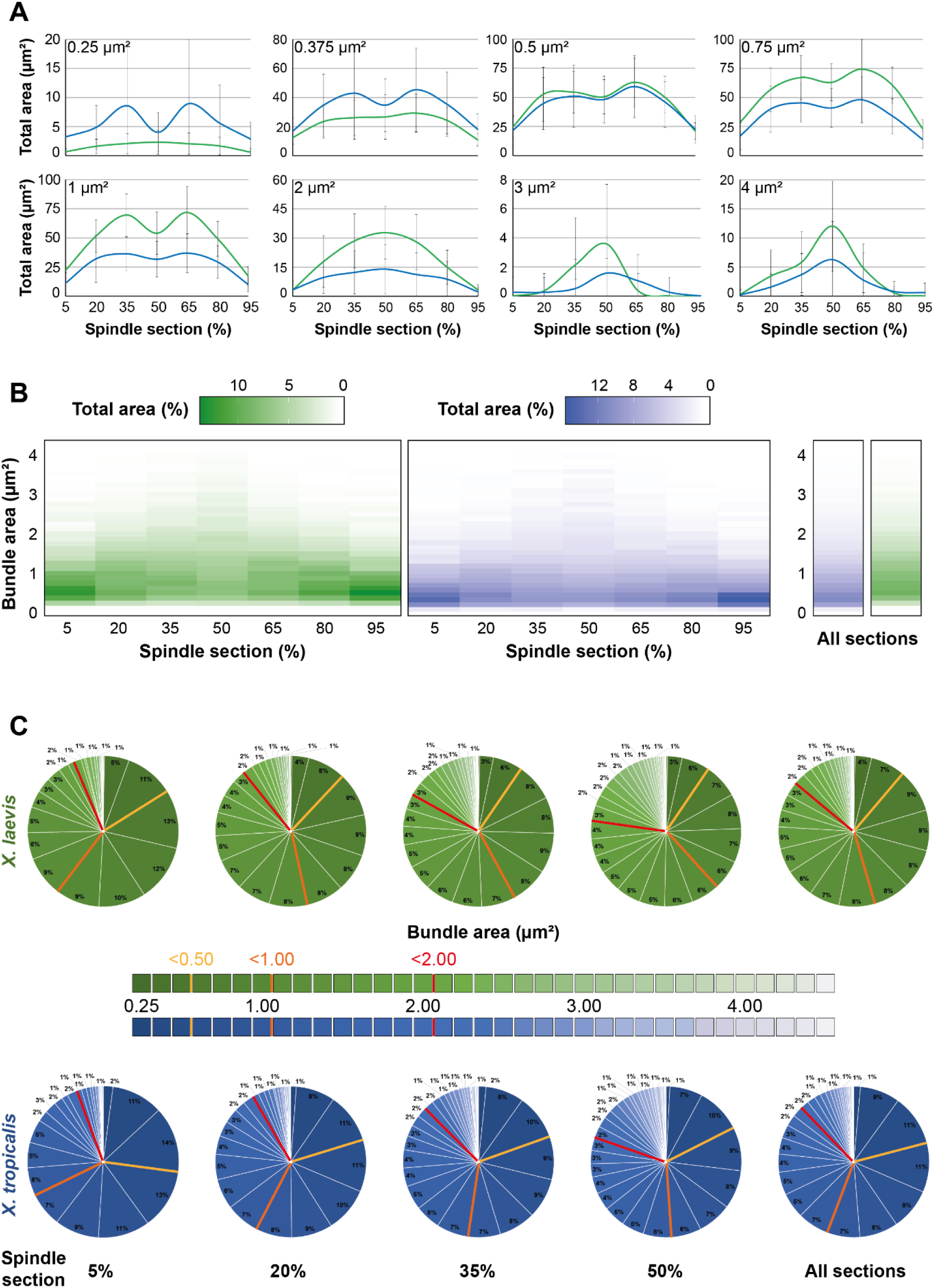
Analysis of the organizational differences between spindles from *X. laevis* and *X. tropicalis*. (A) Normalized average total area of indicated bundle area size bins (top left corners) according to the spindle cross-sections from *X. laevis* (green) and *X. tropicalis* (blue). Note that the absence of left-right symmetry for the distributions of large bundles (3 and 4 µm^2^) is the results of their low count (See Figure 6). (B) Heatmaps representing for each bundle area size bin (y-axis) its normalized average total area (color intensity) for the different indicated cross-sections of the spindle (x-axis) (left) or for all sections together (right) from *X. laevis* (green) and *X. tropicalis* (blue). (C) Pie charts of normalized average total area of bundles according to their binned areas for the different indicated cross-sections of the spindle or for all sections together (right pie charts) from *X. laevis* (green) and *X. tropicalis* (blue). Colored bars in pie charts delimit areas of greater classes of bundle areas (bin 0.0 – 0.5 µm^2^, yellow; bin 0.625 – 1.0 µm^2^, orange; bin 1.125 – 2.0 µm^2^, red). For (A-C), bundle areas are binned into classes of 0.125 µm^2^. Each bin is named according to its upper range value; n = 11 spindles from three egg extracts, n = 15 spindles from three egg extracts, respectively for *X. laevis* and *X. tropicalis*.

We next generated heatmaps of the total areas of each bundle area classes across the spindle sections to provide a clearer overview and compare the space occupied by the different bundle sizes over the spindle length between *X. laevis* and *X. tropicalis* (Figure 7B). We observed that smaller bundles occupied most of the spindle pole area, and the larger bundles took up more space towards the spindle center. Additionally, it became clearly visible that the global area of *X. tropicalis* spindles were predominantly made up of smaller bundles, while *X. laevis* spindles presented a broader range of bundle sizes.

In a different view, pie charts allowed to us to study partitioning of the differently sized microtubules bundles in each section of the spindle (Figure 7C). In the 5% section, very large (2.0 – 4.0 µm^2^), large (1.0 – 2.0 µm^2^), and small (0.5 – 1.0 µm^2^) bundles were divided in similar proportions between *X. laevis* and *X. tropicalis* (7 vs. 4, 33 vs. 26, 44 vs. 40%, respectively) with the main difference being for the very small bundles (0.0 – 0.5 µm^2^), which represented 27% in *X. tropicalis* and only 16% in *X. laevis*. At the 20% section, more than half the section was occupied by large and very large bundles (10 and 43%) in *X. laevis,* whereas this section was divided among very small, small, and large bundles (20, 38 and 32%) in *X. tropicalis*. In the 35% section, the distribution was nearly the same as for the 20% section, except that in both species the proportion of very large bundles nearly doubled. Interestingly, at 50%, there were similar numbers of large bundles and very small bundles in *X. tropicalis* (∼20%), whereas in *X. laevis*, although very large bundles also represented 20%, very small bundles accounted for only half of that in *X. tropicalis*. Finally, pie charts combining all sections summarized these observations. In *X. laevis*, large bundles were prominent, followed by small bundles and an equal amount of very large and very small bundles. In *X. tropicalis*, large and small bundles equally shared the majority, followed by very small bundles, which accounted for twice as much as the very large bundles.

Altogether, these results reveal clear differences in microtubule organization between *X. laevis* and *X. tropicalis.* These differences are likely to be important for building spindles of different sizes while ensuring proper pole stability as well as bipolarity. Further investigation will be necessary to link our observations to the underlying mechanisms that generate distinct spindle architectures.

## DISCUSSION

Electron microscopy techniques can achieve single microtubule resolution, but analyzing entire *Xenopus* spindles remains out of reach due to their large size. Meanwhile, the high microtubule density in these spindles, together with the low resolution of traditional fluorescence microscopy, makes it challenging to accurately study their organization beyond global spindle architecture. As a result, our understanding of microtubule arrangement within *Xenopus* spindles has remained largely speculative. In this study, we developed an optimized, simple and versatile expansion microscopy protocol, coupled with a specifically developed analysis workflow, to address this knowledge gap and compare *X. laevis* and *X. tropicalis* spindles.

Using our analysis workflow, we could reproduce previous observations that tubulin fluorescence intensity is higher at the poles and reduced in the center of the *X. tropicalis* spindles, whereas *X. laevis* spindles exhibit more uniform tubulin distribution (Kitaoka *et al*., 2018) (Supplemental Figure S4C-D). Yet, our detailed analysis revealed distinct microtubule arrangements and variations in spindle architecture that were not detectable with earlier imaging methods. We found that *X. laevis* spindles exhibit a broader range of bundle sizes, while *X. tropicalis* spindles are more limited to smaller bundles (Figure 6E; Figure 7B), with approximately 30% fewer bundles overall. This suggests that *X. tropicalis* could compensate for its fewer microtubules by maintaining smaller bundle sizes while increasing bundle density (Supplemental figure S4C-D). Additionally, we observed that both species favor larger bundle sizes (3-4.5 µm² classes) near and at the spindle center, but *X. tropicalis* spindles prefer very small bundles (0.125-0.375 µm²), while *X. laevis* spindles favor medium-sized bundles (0.75-1.375 µm²) to populate the spindle between the poles and the mid-zone (Figure 7A). This suggests that spindle architecture is regulated to optimize bundle size, which could be crucial for maintaining spindle integrity in smaller spindles, while ensuring chromosome attachment and bipolarity with less flexible bundle sizes. Notably, if the large bundles at the spindle center, of the same sizes between *X. laevis* and *X. tropicalis*, were in part k-fibers, this could explain why k-fibers would need to be specifically differentially regulated by katanin between these two species to maintain spindle integrity in the context of differentially sized spindles (Loughlin *et al*., 2011). Taken together, these data offer new insights to better understanding of how spindle architecture adapts to different morphologies.

In developing this protocol, we made several important technical observations. First, rather than relying on an existing protocol, we reevaluated several steps of classical expansion protocols. This was motivated by the fact that even within the same cell, different cellular organelles can exhibit varying expansion factors (Büttner *et al*., 2021). Additionally, the development of a holed metal slide to image the same spindles before and after expansion proved to be key. This allowed us to calculate the local expansion factor for each spindle rather than relying on the global gel expansion (Truckenbrodt *et al*., 2019), and systematically assess the quality of expansion under different conditions. Notably, we found that the most critical step was actually sample fixation. Both aldehydes fixatives and dehydration fixation can introduce artifacts (Cross and Williams Jr., 1991; Melan, 1999). We observed that fixation causes spindle shrinkage and correlates with wavy microtubules (Figure 5E). In addition, over-fixation led to poor incorporation of structures into the gel and disturbed expansion (Supplemental Figure S2D), while under-fixation caused spindles to disappear between pre- and post-expansion imaging, likely because they washed off during processing. We also aimed to preserve the rhodamine-labeled tubulin signal and adequately fix the sample onto the coverslip for pre-expansion immunolabeling (Supplemental Figure S3). It is therefore only by carefully and systematically evaluating the different parameters and the resulting expansion quality, and finding the proper balance that we were able to develop this optimized protocol.

Through this study, we established a simple ExM protocol optimized for the *Xenopus* egg extract system. This protocol provided new insights into spindle architecture adaptation and will facilitate further high-resolution studies of spindle components or other microtubule structures, such as asters, as well as spindles from other *Xenopus* species (Kitaoka *et al*., 2018; Miller *et al*., 2019).

## MATERIALS AND METHODS

A Bio-protocol manuscript of our optimized fixation and expansion protocol is written and ready to be submitted.

### Animals

All animal experimentation in this study was performed according to our animal use protocol APAFiS #26858-2020072110205978 approved by the Animal Use Ethic Committee (#7, Rennes, France) and the French Ministry of Higher Education, Research and Innovation. Mature *X. laevis* and *X. tropicalis* female frogs were ovulated with no harm to the animals with rest intervals between ovulations of at least a 6-month and 3-month, respectively.

### Chemicals

Unless otherwise stated, all chemicals were purchased from Sigma-Aldrich, Merck.

### *Xenopus* egg extracts

*X. laevis* and *X. tropicalis* egg extracts were prepared and cycled spindle reactions conducted as previously described (Maresca and Heald, 2006; Kitaoka *et al*., 2024). Briefly, eggs arrested in metaphase of meiosis II were collected, dejellied, and fractionated by centrifugation. The cytoplasmic layer was isolated, supplemented with 10 mg/mL each of leupeptin, pepstatin, and chymostatin (LPC) protease inhibitors, 20 mM of cytochalasin B, a creatine phosphate and ATP energy regeneration mix, as well as 2 µM rhodamine-labeled porcine brain tubulin (Cytoskeleton). Then, cycled spindle reactions containing 25 µL of cytostatic factor-arrested (CSF) extract from *X. laevis* or *X. tropicali*s were supplemented with 400 µM Ca2+ and incubated for 5.5 min at 23°C to bring the extract into interphase. Resulting interphasic extract was supplemented with *X. laevis* or *X. tropicali*s sperm nuclei, prepared as previously published (Hazel and Gatlin, 2018), at a final concentration of 500 to 1,000 per µL. After about 45 min (30 min for *X. tropicalis*), the DNA was replicated and 50 % of fresh CSF extract was added to the reaction leading to cycle the extract back to metaphase, which then produced bipolar spindles around 45 min after adding back at 23°C.

### Spindle spindown

When organized bipolar spindles were observed, the ∼50 µL reactions were mixed to 1 mL of dilution solution containing 30% glycerol, 1X BRB80 (BRB80 5X: 400 mM 1,4-Pipperazinediethanesulfonic acid (PIPES), 5 mM MgCl2, 5 mM EGTA, pH 6.8), 0.5% Triton X-100 and supplemented with 2.5% or 5% formaldehyde or 0.05%, 0.10%, 0.15% or 0.25% glutaraldehyde (Electron Microscopy Science), or no fixative and layered over a 5 mL cushion containing 40% glycerol in 1X BRB80. The reactions were then spun onto coverslips at 17,000 g for 15 min or 6,000 g for 20 min at 16°C. After that, the samples were fixed on the coverslips by putting them in −20°C methanol, Dent’s (Methanol / DMSO, 4:1), MAD (Methanol / Acetone / DMSO, 2:2:1) or MAD2 (Methanol / Acetone / DMSO, 3:1:1) baths for 2 s to 5 min depending on the protocol. Coverslips were then put on parafilm with the sample facing up in a large petri dish kept humid with wet kimwipes and washed trice with PBS supplemented with 0.1% (v/v) IGEPAL® CA-630 (PBSI) to stop dehydration and rehydrate the samples and then twice with PBS. Coverslips were finally processed either for pre-gelation immunofluorescence, pre-expansion imaging or directly for anchoring.

### Pre-expansion imaging

For pre-expansion imaging, coverslips were first incubated for 5 min with PBS containing 10 µg/mL of Hoechst 33342 and rinsed twice with PBS. Coverslips were then mounted on the cavity of a homemade metal slide filled with PBS. Imaging was performed on an Olympus BX51 microscope equipped with a Lumencor SOLA SE U-nIR light source, and a Photometrics Prime-BSI sCMOS Back Illuminated camera. Images were acquired using the μManager acquisition software v1.4 (Edelstein *et al*., 2014). After imaging, an excess of PBS was added into the cavity of the metal slide to retrieve the coverslip and replace it on the parafilm of the humid petri dish shielded from light and recovered with PBS.

### Sample anchoring and gelation

The PBS was removed, the coverslips were rinsed once with 120 µL of anchoring solution (0.1 mg/ml 6-((Acryloyl) amino) hexanoic acid, succinimidyl ester (AcX,SE; ThermoFischer Scientific)), and incubated in 120 µL of fresh anchoring solution for 1h at room temperature and then processed for gelation. A drop of 75 µL of ExM gelation mix (Chen *et al*., 2015; Tillberg *et al*., 2016) (8.625% (wt/wt) Sodium Acrylate, 2.5% (wt/wt) Acrylamide, 0.15% (wt/wt; Interchim) N,N’-methylenebisacrylamide, 2 M NaCl in 1X PBS) or U-ExM gelation mix (Gambarotto *et al*., 2018) (19% (wt/wt) Sodium Acrylate, 10% (wt/wt) Acrylamide, 0.1% (wt/wt) N,N’-methylenebisacrylamide in 1X PBS) supplemented with 0.2 % TEMED and then 0.2% Ammonium persulfate was placed on the parafilm. After aspiring all the anchoring solution, the coverslips were rapidly but carefully put on the drop with the spindles facing the gelation solution. After 10 min of polymerization at room temperature, the wet kimwipes were removed and the reaction was then left to polymerized for 30 min at 37°C in the petri dish shielded from light.

### Digestion or denaturation of samples in the gels

For the digestion procedure, the coverslips and their attached gel were transferred in a small glass petri dish containing the digestion buffer (1X TAE, 0.5 % Triton X-100, 0.8M guanidine hydrochloride). Coverslips were gently removed from the gel with a spatula and the buffer was supplemented with proteinase K (New England Biolabs) at 8 U / mL at the last moment. The digestion was performed at 37°C for 1h. Gels were then collected with the spatula and put in new small petri dish, washed 3 times for 5 min in PBS supplemented with 0.1% Triton X-100 (PBT), twice in PBS and processed for expansion.

For the denaturation procedure, the coverslips and their attached gel were transferred in a small glass petri dish and incubated for 15 min at room temperature in denaturation buffer (200 mM SDS, 200 mM NaCl and 50 mM Tris (Euromedex), pH 9). Coverslips were then gently removed from the gel with a spatula and gels were incubated in fresh denaturation buffer for 30 min in a water bath at 95°C. Following denaturation, the gels were washed 3 times for 5 min in PBT, twice in PBS and processed either for post-gelation immunofluorescence or expansion.

### Expansion

Gels, in glass petri dish, were first incubated for 5 min at room temperature with 1 mL of the Hoechst solution diluted in PBS. Following 3 washes with PBS, MilliQ water was added and left for 1 h. Gels were further expanded by changing for fresh milliQ water and left for at least 30 min before imaging. After imaging, gels were put back into 1X PBS, rinsed once and stored at 4°C shielded from light to preserve the fluorescent signal.

### Post-expansion imaging

To compare the same spindles before and after expansion, the whole gel was mounted on two microscope slides taped together. The entire gel can then be screened, keeping its integrity and allowing to retrieve all structures of interest. To mount the gel, the milliQ water was removed, the gel taken out and put on a kimwipes to dry it. The gel was then transferred to the double slide and covered with two large (24x60) coverslips to be imaged on the Olympus BX51 microscope as for pre-expansion imaging.

For greater resolution imaging, a Zeiss LSM 880 confocal microscope with Airyscan was used. Expanded gels were cut into small pieces to be mounted on a glass bottom dish (MatTek) coated with poly-L-lysine to prevent gel drifting during imaging. For a better gel adhesion to the glass, gel pieces were dried on kimwipes before mounting. The inverted Zeiss Axio Observer 7 microscope equipped with Confocal Scan Head LSM 880 and Airyscan detector was used in Fast-Airyscan mode and images were acquired using a 40x/1.2 W C Apochromat objective with powers of 20% and 30% for the 561 nm and 405 nm lasers, respectively, and spacing of 0.25 µm in Z. Images are mean averages of 2 scans with a depth of 16 bits. Channels were acquired sequentially, and pinhole size was always chosen to correspond to 1 airy unit.

### Optional pre-gelation immunofluorescence

For our proof of principle, coverslips were first blocked with PBS supplemented with 3% BSA (Euromedex) (PBS-BSA) overnight at 4°C in a large humid petri dish. On the next morning, each coverslip was washed with PBS-BSA. Then 110 µL of PBS-BSA containing 1:250 of anti-dynein mouse antibody (Invitrogen, #14-9772-80) was put on each coverslip for 1h at room temperature in the large humid petri dish. Coverslips were then washed with PBSI 3 times for 5 min and subsequently incubated for 30 min at room temperature with 110 µL of PBSI containing 1:500 of goat anti-mouse AlexaFluor488-coupled antibody (Invitrogen, # A-11029) in the large humid petri dish shielded from light. Coverslips were washed in PBSI 3 times for 5 min and then incubated for 5 min at room temperature with 110 µL of the Hoechst solution diluted in PBSI. Coverslips were washed in PBSI 3 times for 5 min and twice with PBS. Coverslips were then processed either for pre-expansion imaging and anchoring.

### Analysis of spindle parameters

Length and normalized intensity distribution profile were measured for spindles. For each spindle length a line with a width of 10 pixels for unexpended and 40 pixels for expanded spindles was drown from one pole to another in Fiji (Schindelin *et al*., 2012). Intensity line scans measured from pole to pole were then normalized to 100% of the length of each spindle using an automated Java ImageJ plugin developed by Dr. Xiao Zhou (https://github.com/XiaoMutt/AiSpindle). The intensities were then also normalized to the maximum intensity of each dataset. To compare tubulin distribution between pre- vs. post-expansion, the same spindles were systematically analyzed in the same order and with the line carefully placed at the same location and in the same orientation.

Expansion factors and distortions were measured using Elastix (http://elastix.isi.uu.nl) and a Mathematica (Wolfram) script as previously described (Chozinski *et al*., 2016). Briefly, post-expansion images were aligned with corresponding pre-expansion images using linear transformations (rotation, scaling, and translation). The linearly transformed post-expansion images were then nonlinearly deformed to match the pre-expansion images. Quantitative comparison of the linearly and nonlinearly transformed post-expansion images produced values for root-mean-square (RMS) deviation over a range of length scales. Each RMS plot was generated with one set of representative pre- and post-expansion images.

Curvature measures were performed on Fiji using Kappa plugin (Mary and Brouhard, 2019) following microtubule bundles from pole-to-pole on expanded spindles.

### Expanded spindle bundle analysis workflow

#### Code

The workflow was coded as an ImageJ (Schneider *et al*., 2012) macro and is available at “https://github.com/tpecot/IJmacro_MicrotubuleBundlesQuantification/blob/main/MicrotubulesBundleQuantification.ijm”. The Stardist model trained to segment bundles of microtubules is available at “https://github.com/tpecot/IJmacro_MicrotubuleBundlesQuantification/blob/main/TF_SavedModel.zip”.

#### Spindle poles selection

The user manually defines the location of the two spindle poles to accurately localize those two reference points.

#### Background subtraction

An optional top-hat transform can be applied to the image to subtract background with CLIJ plugin (Haase *et al*., 2020b, 2020a). A scaling is then performed to obtain an image volume with isotropic voxel resolution. This option was not used in our analysis.

#### First registration

From the coordinates of spindle poles, a translation in the xy plane is first applied to position the spindle center at the image center and a rotation around z axis is performed to align the spindle axis with the x axis. Those two transformations are implemented with the CLIJ suite plugin (Haase *et al*., 2020b, 2020a).

#### Getting rid of intensities coming from microtubules of other spindles

As other spindles might be close and acquired in the same image, it is possible to define the spindle area with a polyline ROI to exclude intensity coming from outside the spindle of interest.

#### Second registration

From the coordinates of the spindle poles, a rotation around the y axis is applied to align the spindle axis along the z axis with the TransformJ plugin (Meijering *et al*., 2001). The registered volume shows now cross sections of microtubule bundles.

#### Bundle segmentation

Microtubule bundles at selected cross sections (possible from 5% to 95%, every 5%) are segmented with the deep learning approach Stardist (Schmidt *et al*., 2018). Although Stardist was designed for nuclei segmentation, the available pre-trained models did surprisingly okay on segmenting the bundles but did not show a sufficient accuracy performance. A new model was therefore trained on 10 images showing cross sections of microtubules, of which 3,162 were manually annotated, 2 images were used for validation. For training, images were normalized with a 1-99.8 quantile. The new model was initialized with the versatile fluorescence model and trained for 1,000 epochs. The trained model was converted as a TF_Saved model to be used in the Stardist Fiji plugin.

#### Measurements and result visualization

A list of measurements (centroid coordinates, area, perimeter, ellipse fitting, average intensity, total intensity) were exported for each bundle and combined in an Excel file with the Excel function plugin. At each position, the image of bundles and the segmentations displayed as outlines is also generated to visually inspect the results.

### Image visualization, data compilation and statistics

Image stacks were visualized in Fiji (ImageJ 1.54f) and in Napari (version 0.4.19; napari.org) and processed in Fiji. Data were analyzed and statistics performed using Excel (Microsoft), Mathematica (Wolfram) and the R suite (www.r-project.org). Averages and standard deviations were calculated. One-way ANOVA with post hoc Dunett’s test were performed to test for statistical significance.

## ACKNOWLEDGMENTS

We thank the members of the Gibeaux and the Heald labs for their support. We thank the members of the FAIIA and MRic facilities, in particular Dr. Xavier Pinson. We are grateful to Paul Guichard and Virginie Hamel for guidance in developing this protocol as well as Julien Maurais for taking great care of our frogs. G.G. was supported by PhD fellowships from the Region Bretagne (ARED) and from the Fondation pour la Recherche Médicale (FRM) FDT202304016369, and M.K. by a National Science Foundation (NSF) GRFP fellowship. R.H. was supported by NIH MIRA grant R35 GM118183 and the Flora Lamson Hewlett Chair, and R.G. by a Human Frontier Science Program grant CDA00019/2019-C.

## ABBREVIATIONS USED

ExM: expansion microscopy
AcX: succinimidyl ester of 6- ((acryloyl) amino) hexanoic acid
U-ExM: ultrastructure expansion microscopy
RMS: root-mean-square
ROI: region of interest
PBSI: PBS supplemented with 0.1% (v/v) IGEPAL® CA-630.

---

**SUPPLEMENTAL FIGURE S1:**
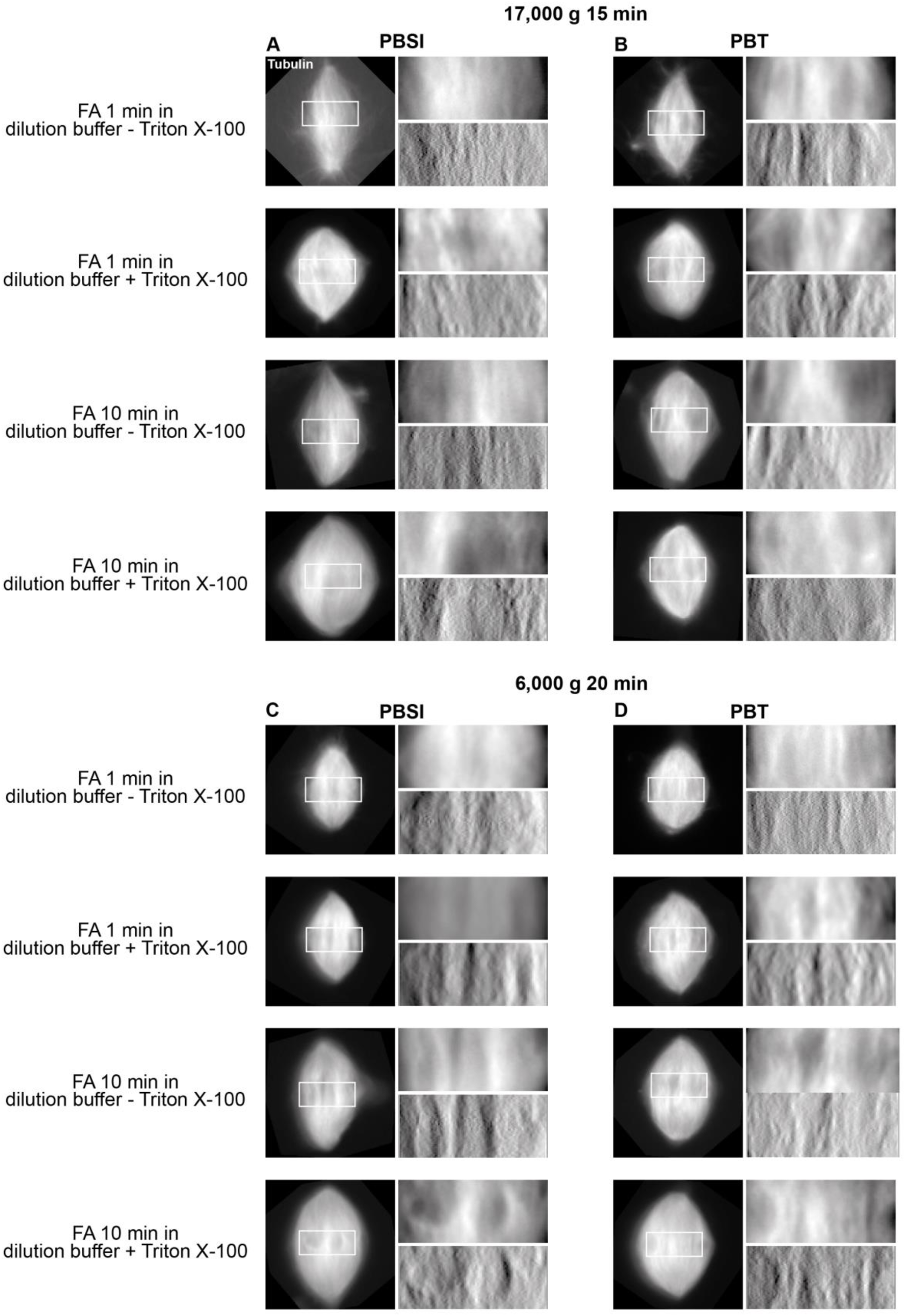
Evaluating the effects of varying different fixation and spindown conditions. Samples were spun-down at 17,000 g during 15 min (A-B), or at 6,000 g during 20 min (C-D), and then rehydrated with PBSI (A and C), or PBT (B and D). For each, the condition of fixation is indicated on the left. For each, top right images show magnified views of boxed regions in left images, representing unexpanded spindles after spin-down, with rhodamine-labeled tubulin signal shown. Bottom right images are top right images processed using the Gradient-X filter from the OrientationJ plugin to enhance the visualization of microtubules, highlighting their orientation in Y and improving the contrast. Scale bars, 10 µm; PBSI: PBS + 0.1% IGEPAL® CA-630; PBT: PBS + 0.1% Triton X-100; FA: Formaldehyde.

**SUPPLEMENTAL FIGURE S2:**
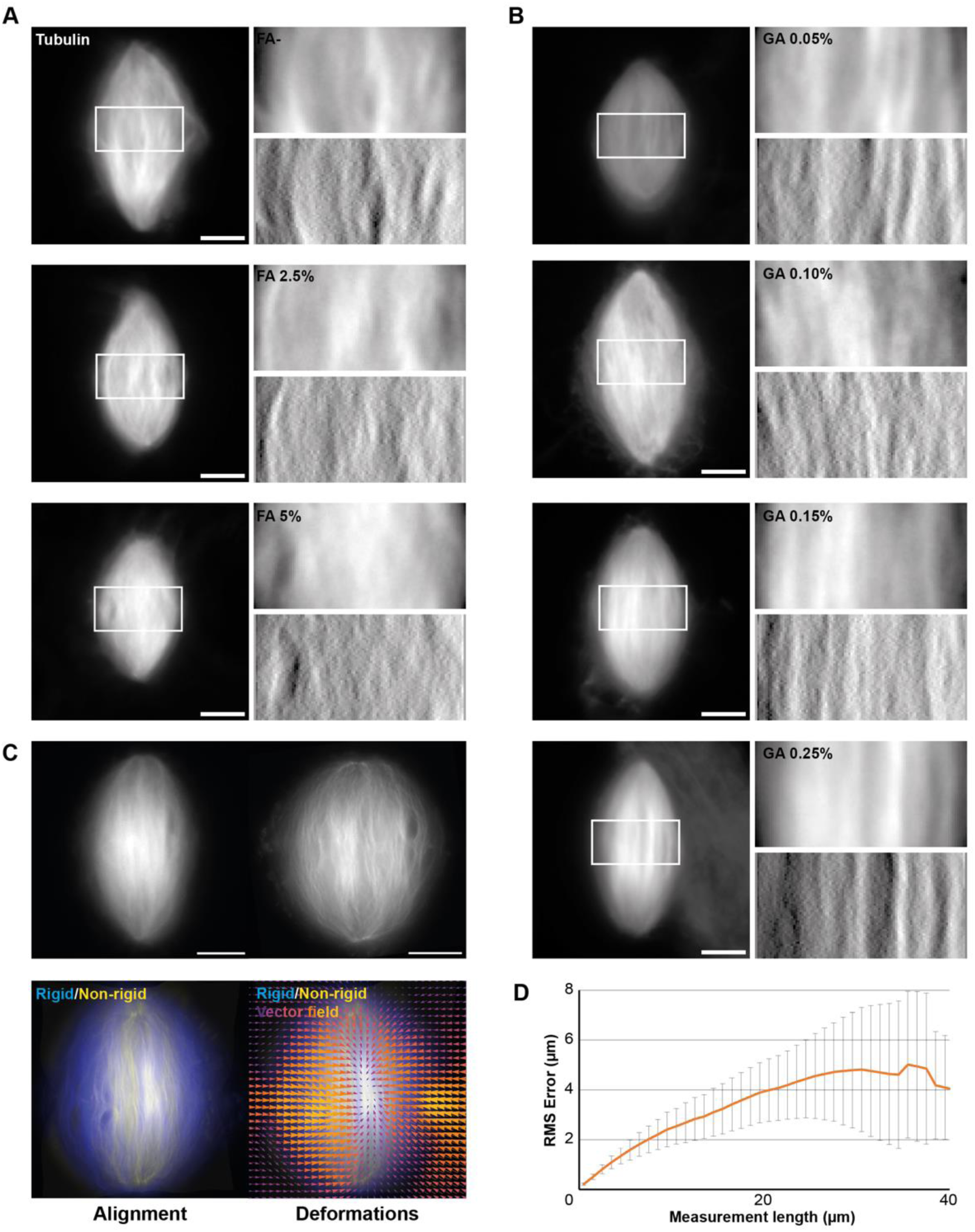
Evaluating the effects of changing pre-centrifugation chemical fixation. Samples were fixed with formaldehyde (A) or glutaraldehyde (B). For each, left images are unexpanded spindles fixed with indicated concentration of fixators with rhodamine-labeled tubulin signal shown. Top right images are magnified views of boxed regions in corresponding left images. Bottom right images are top right images processed using the Gradient-X filter from the OrientationJ plugin to enhance the visualization of microtubules, highlighting their orientation in Y and improving the contrast. (C) Top images are same unexpanded (left) and expanded (right) spindle fixed with 0.25% glutaraldehyde. Bottom left image show the alignment made between rigid registration and non-rigid registration, and bottom right, the vector field of deformations calculated from the registrations. (D) Root-Mean-Square length measurement error as a function of measurement length for pre- vs. post-expansion images. The orange line shows the average and the error bars, the standard deviation (n = 16 spindles from one egg extract). Scale bars, 10 μm for unexpanded spindles and 40 µm for expanded spindles.

**SUPPLEMENTAL FIGURE S3.**
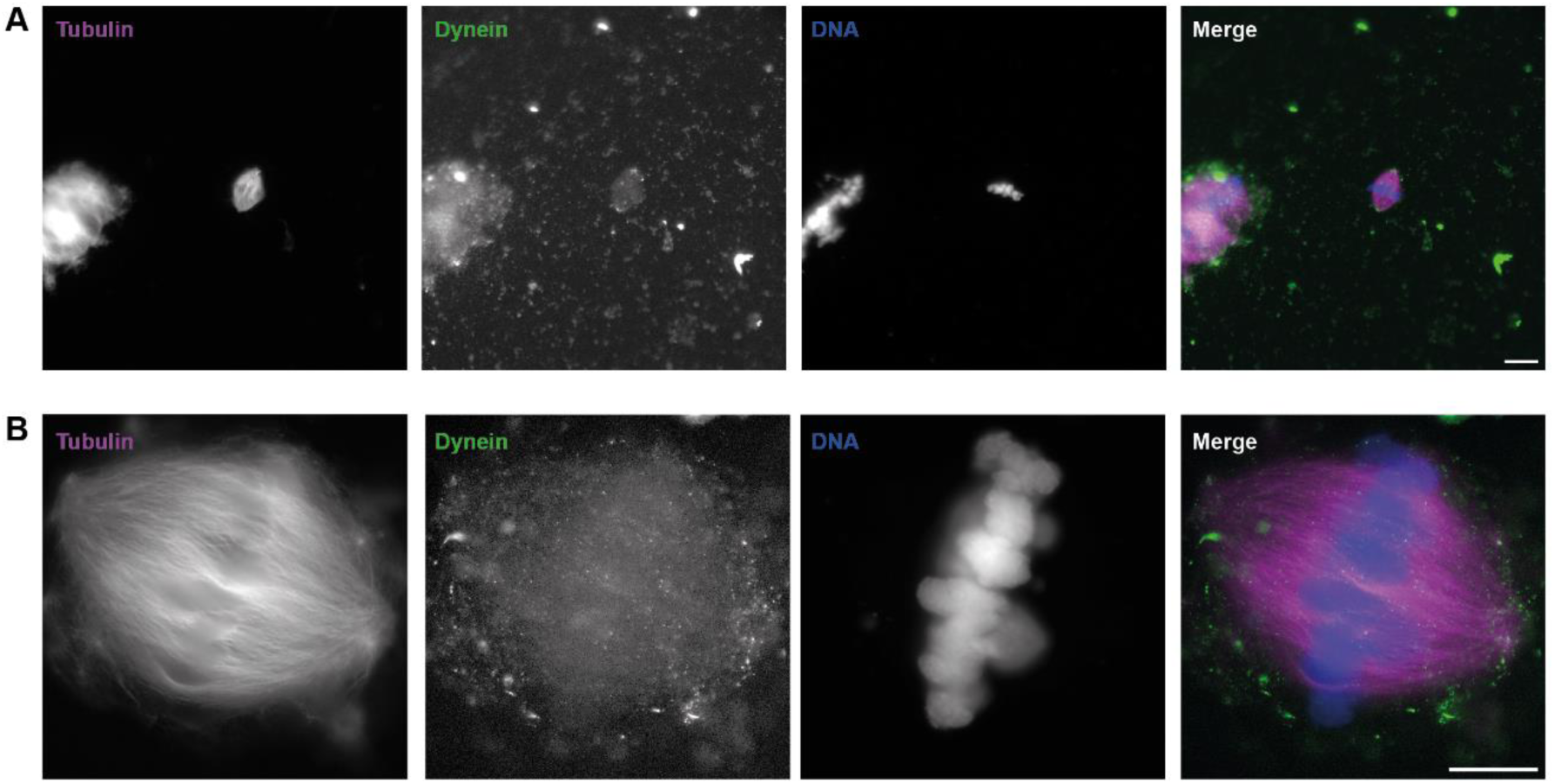
Combining pre-expansion immunolabeling and expansion microscopy. Unexpanded (A) and expanded (B) spindles fixed for 10 s in cold methanol bath and immunolabeled with mouse anti-dynein primary antibody, and goat anti-mouse AlexaFluor488-coupled secondary antibody. Note that the two spindles presented here are different ones, as we did not aim for pre- and post-expansion correlation for this analysis. For each, tubulin signal is shown in left images, dynein signal in middle left images, DNA signal in middle right images, and a merged channels view is shown in right images. Scale bars, 20 µm for unexpanded spindle, 40 µm for expanded spindle.

**SUPPLEMENTAL FIGURE S4:**
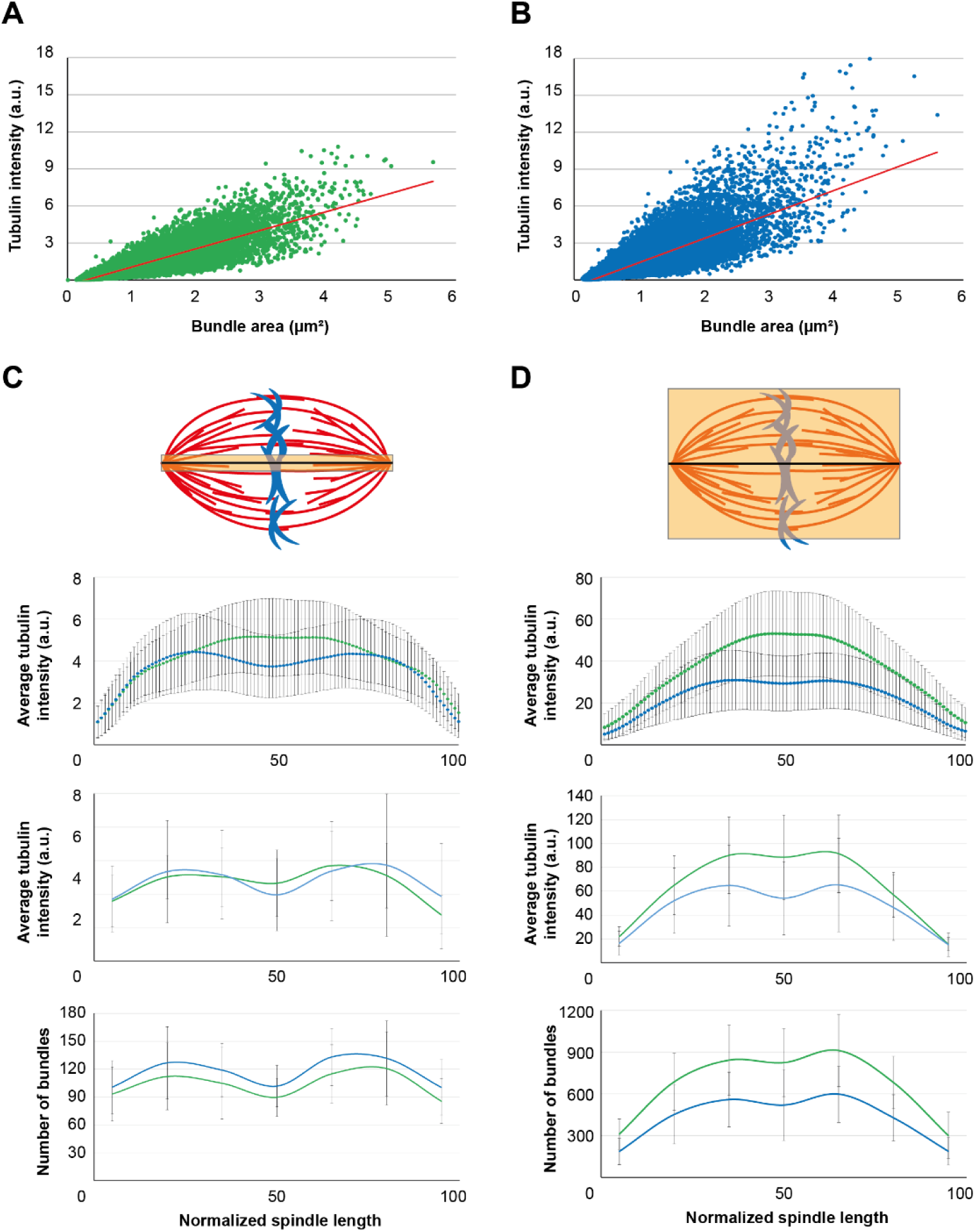
Assessing the accuracy of microtubule bundle segmentation. (A) Representation of area for all detected microtubule bundles in *X. laevis* spindles as a function of their fluorescence intensity, with the trend line in red (R^2^ = 0.76, n = 53,058 bundles from 11 spindles from three egg extracts). (B) Representation of area for all detected microtubule bundles in *X. tropicalis* spindles as a function of their fluorescence intensity, with the trend line in red (R^2^ = 0.69, n = 54,880 bundles from 15 spindles from three egg extracts). (C) Top sketch representing a spindle analyzed with a small line scan. Top panel is the average fluorescence intensity distribution ± standard deviation along thin-line-scanned *X. laevis* (green) and *X. tropicalis* (blue) unexpanded spindles. Middle panel is the average fluorescence intensity distribution ± standard deviation along thin-ROI-analyzed *X. laevis* (green) and *X. tropicalis* (blue) expanded spindles. Bottom panel is the average number of detected bundles along *X. laevis* (green) and *X. tropicalis* (blue) expanded bundles. (D) Top sketch representing a spindle analyzed with a line scan entirely covering the spindle. Top panel is the average fluorescence intensity distribution ± standard deviation along thick-line-scanned *X. laevis* (green) and *X. tropicalis* (blue) unexpanded spindles. Middle panel is the average fluorescence intensity distribution ± standard deviation along fully analyzed *X. laevis* (green) and *X. tropicalis* (blue) expanded spindles. Bottom panel is the average number of all detected bundles along *X. laevis* (green) and *X. tropicalis* (blue) expanded bundles. For (C) and (D), n = 59 unexpanded *X. laevis* spindles; n = 53 unexpanded *X. tropicalis* spindles; n = 11 expanded *X. laevis* spindles; n = 15 expanded *X. tropicalis* spindles; all from three independent egg extracts.

**SUPPLEMENTAL MOVIE S1:** *X. laevis* egg extract spindle expanded with our optimized fixation and expansion protocol and visualized in Napari. Z-stacks were acquired on an inverted Zeiss Axio Observer 7 microscope equipped with Confocal Scan Head LSM 880 and Airyscan detector used in Fast-Airyscan mode and images were acquired with a depth of 16 bits using a 40x/1.2 W C Apochromat objective with powers of 20% of the 561 nm laser, a spacing of 0.25 µm in Z and pinhole size corresponding to 1 airy unit. The tubulin signal is shown in gray.

**SUPPLEMENTAL MOVIE S2:** *X. tropicalis* egg extract spindle expanded with our optimized fixation and expansion protocol and visualized in Napari. Z-stacks were acquired on an inverted Zeiss Axio Observer 7 microscope equipped with Confocal Scan Head LSM 880 and Airyscan detector used in Fast-Airyscan mode and images were acquired with a depth of 16 bits using a 40x/1.2 W C Apochromat objective with powers of 20% of the 561 nm laser, a spacing of 0.25 µm in Z and pinhole size corresponding to 1 airy unit. The tubulin signal is shown in gray.

## Notes

### Competing Interest Statement

The authors have declared no competing interest.

### Summary of Updates

Minor corrections were made throughout the manuscript, including typos, grammar, and phrasing for clarity. A few references were also updated.

